# Chemical circularization of in vitro transcribed RNA opens new avenues for circular mRNA design

**DOI:** 10.1101/2024.10.10.617555

**Authors:** Adam Mamot, Malgorzata Wasinska-Kalwa, Karol Czubak, Katarzyna Frankowska, Tomasz Spiewla, Marcin Warminski, Dominika Nowis, Jakub Golab, Joanna Kowalska, Jacek Jemielity

## Abstract

Circularization is at the frontier of therapeutic messenger RNA (mRNA) enhancements. Currently available enzymatic and ribozymatic methods for generating circular RNAs (circRNAs) face several challenges related to sequence limitations, purification, and sub-optimal biological activity. The chemical circularization of synthetic RNA fragments potentially overcomes these limitations but is applicable only to extremely short sequences. Here, we report a novel approach for accessing circular RNAs based on the chemical circularization of in vitro transcribed RNA. We efficiently accessed chemically circularized RNAs (chem-circRNAs) by making in vitro transcribed precursor RNAs modified at the 5′ end with an ethylenediamine moiety, which undergoes an intramolecular reaction with the periodate-oxidized RNA 3′ end under reductive amination conditions. We demonstrate that this method is modification-compatible and applicable to various sequences. Additionally, we report methods for the effective separation of chem-circRNAs from their linear precursors. Using this approach, we prepared multiple chemically-obtained circular RNAs (chem-circRNAs; 35–1500 nt long) with circularization efficiencies reaching up to 60%. We show that protein-coding chem-circRNAs are translationally active in living cells and exhibit increased durability, similar to enzymatically circularized mRNAs. We also demonstrate that this approach enables unprecedented access to chemically modified circRNAs, such as circ-mRNAs incorporating a functional endocyclic N7-methylguanosine cap or modified with N1-methylpseudouridine within the RNA body. Notably, circRNAs containing an endocyclic cap structure engage in the most efficient, cap-dependent mechanism of translation. Our approach makes chemically-modified circularized full-length protein-coding RNAs easily accessible, thereby opening new avenues for the design, modification, and functionalization of circular mRNAs.

## INTRODUCTION

Despite its inherent fragility, mRNA is one of the most prolific substances in the context of therapeutic gene delivery.^1^ To ensure sufficient biological activity of therapeutic mRNA, it is necessary to protect it from premature cleavage by numerous RNA-targeting enzymes, particularly exonucleases.^2^ Circularization is one of the most promising modifications of mRNA in this context, as it prevents exonucleolytic digestion.^3,4^ Consequently, circRNAs have shown higher stability than corresponding linear RNAs and exhibited significantly longer protein expression profiles.^5,6,7^ Unfortunately, the circularization of macromolecular RNA still poses a significant challenge.

Currently established methods of RNA circularization are based on chemical reactions, enzymatic reactions, or the activity of autocatalytic RNA sequences.^8^ The chemical circularization methods rely on the use of short precursor RNA sequences, synthesized via phosphoramidite chemistry.^9-11^ In consequence, the so-far chemically synthetized circRNAs encoded relatively short sequences and have limited use in the context of therapeutic gene delivery. RNA circularization can alternatively be carried out with ligases, such as T4 RNA ligase I and T4 RNA ligase II (Fig. 1A).^12-14^ Both enzymes require 5′ monophosphorylated precursors that are first adenylated and subsequently ligated with the 3′ OH of the ribose at the 3′ end RNA. The ligases have been found to be strongly dependent on target RNA length and sequence.^5,6,14,15^ As a result, it is difficult to predict or improve the performance of such enzymes. Other methods of RNA circularization involve catalytic nucleic acids (Fig. 1A).^5,6,16-19^ Wesselhoeft *et al*. have circularized RNA in vitro by using the permuted intron-exon (PIE) system and optimized self-splicing intronic sequences.^5,6^ Litke *et al*. have shown the Twister-optimized RNA durable overexpression (Tornado) system for RNA circularization in cells.^16^ These methods are based on sequence-dependent activity and can be hindered by the presence of methylated nucleobases [e.g. N1-methylpseudouridine (m^1^Ψ) or N6-methyladenosine] and other chemical modifications.^6^ Moreover, the translation of currently known circRNAs relies on internal ribosome entry site (IRES) sequences,^20,21^ which are less efficient than the main pathways of eukaryotic translation involving cap-and-polyA-dependent initiation.^22-25^ To our knowledge, none of the currently available methods of RNA circularization are compatible with mRNA body modifications or cap-dependent translation mechanisms.

**Figure 1.**
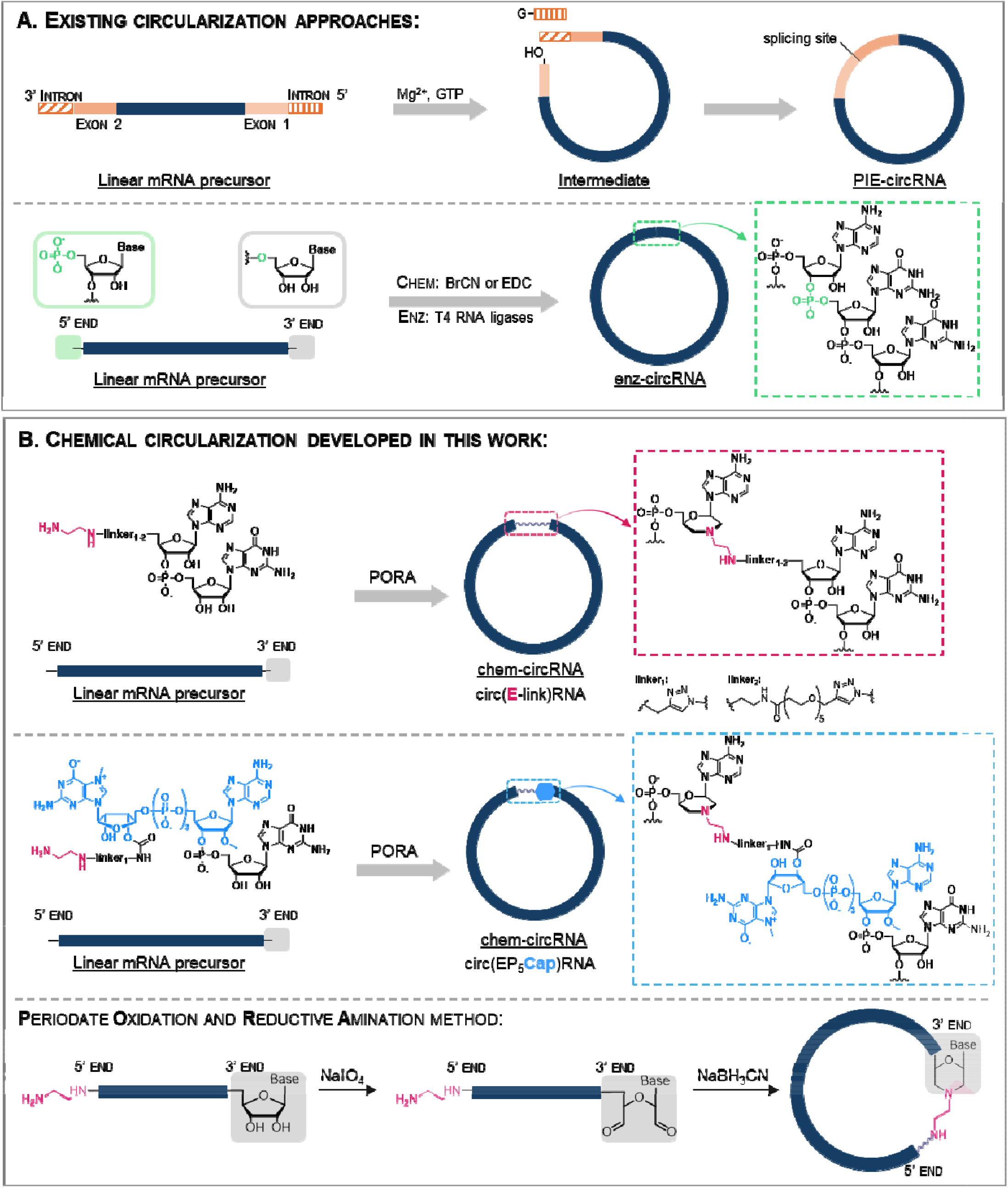
Strategies for RNA circularization: A) Overview of existing circularization approaches. B) Chemical circularization developed in this work. The 5′ modified linear RNA precursor is obtained from a 5′ initiator (EDA-AG, EDA-PEG_5_-AG or EP_5_Cap) during an in vitro transcription (IVT) reaction and is subsequently circularized by PORA reaction. The produced circRNA contains a single chemically modified linkage.

In this work, we developed a chemical circularization method that enables the expansion of circ-mRNA design strategies beyond IRES-dependent translation. Taking advantage of the unique reactivity of ethylenediamine moiety towards oxidized RNA 3′ ends, we developed a post-transcriptional circularization protocol relying on a one-pot, two-step chemical reaction (Fig. 1B).^26^ The protocol is based on affordable, readily available, non-invasive reagents and can be applied to RNA of virtually any size and sequence (Fig. 1B). This approach, combined with improved purification and isolation techniques, provided access to chem-circRNAs of 1500 nt and presumably even longer. mRNAs generated using our approach were evaluated in living cells to determine their relative stability and translational activity. We also show that the novel circularization method is compatible with the incorporation of chemical modifications that were not accessible using currently available approaches – N7-methylguanosine cap and m^1^Ψ. Our findings offer an attractive alternative to chemoenzymatic methods of RNA circularization and open new avenues for circ-mRNA optimization in the future.

## RESULTS

### Development of the chemical circularization method

Our approach towards circRNAs was designed to minimize the number of steps necessary to functionalize and circularize the target RNA, maintaining the process as close as possible to the existing mRNA production pipelines. In the first step, we generate an in vitro transcribed (IVT) pre-circRNA that is 5′ functionalized with ethylenediamine by incorporating a properly designed transcription initiator (primer; Fig. 1B). Such pre-circRNA is then post-transcriptionally circularized by one-pot periodate oxidation and reductive amination (PORA) reaction, resulting in the formation of a morpholine-derived inter-nucleotide linkage (Fig. 1B). As transcription initiators, we designed two types of AG dinucleotides equipped with ethylenediamine linkers (EDA-linkers) of different lengths. We also designed an EDA-functionalized cap 1 analog to provide unprecedented access to circular mRNAs containing an endocyclic 5′ cap structure (Fig. 1B). The presence of circular RNAs was verified by polyacrylamide gel electrophoresis (PAGE) and compared to analogous circRNA obtained by enzymatic ligation (Fig. S2).

The synthesis of the primers was performed in a single synthetic step using click chemistry (NHS-based amidation and CuAAC) from an azido-modified dinucleotide synthesized on solid-support (N_3_-AG, N_3_-A_m_G, Fig. S1).

We began our attempts at chemical RNA circularization using two RNA oligonucleotide models: E-RNA_01_ and E-RNA_02_, (35 nucleotide-long sequences, Table 1, Fig. 2), transcribed in the presence of EDA-AG initiator (Fig. 1B, Fig. S1). The subsequent step of chemical circularization was based on our previous findings that oxidized RNA 3′ ends show particularly high reactivity in PORA reactions toward compounds containing ethylenediamine.^26^ To execute the PORA reaction, E-RNA_01_ and E-RNA_02_ were incubated in a sodium periodate (NaIO_4_) solution, leading to the formation of an acyclic 2′,3′-dialdehyde at the 3′ terminal nucleotide. This derivative spontaneously reacted with ethylenediamine motif to form a cyclic intermediate (Fig. 1B). The subsequent addition of sodium cyanoborohydride (NaBH_3_CN) led to the reduction of the cyclic intermediate, ensuring the irreversibility of the reaction. As a result, a morpholine linkage was formed between the former 3′ terminal nucleotide and the 5′ terminus of the transcription initiator.

**Table 1.**
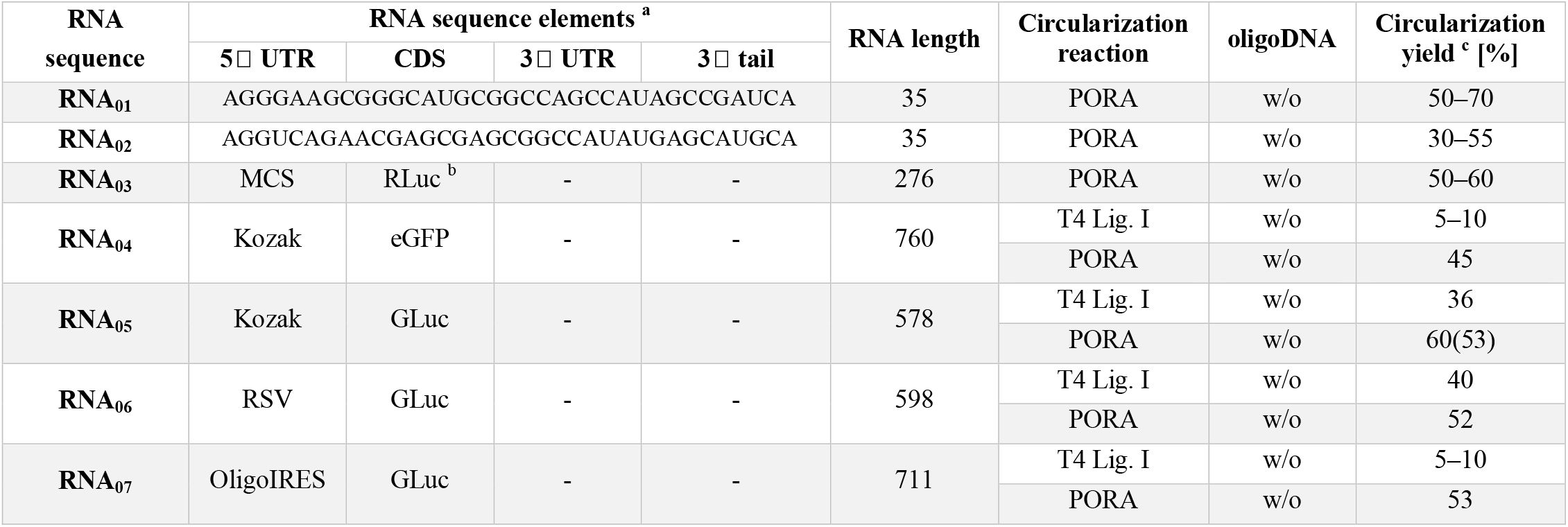
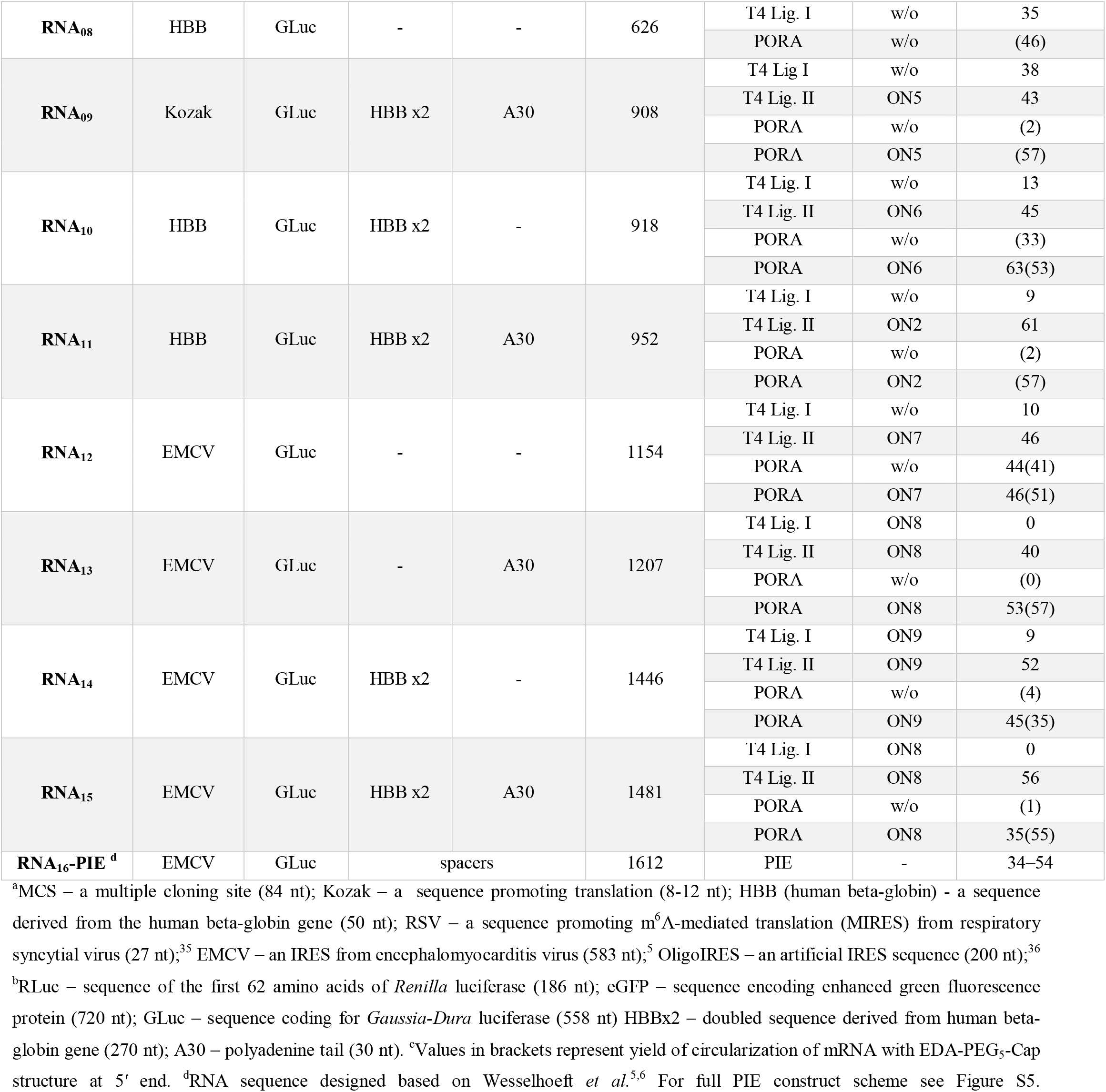
Summary of precursor RNA sequences (abbreviation, schematic of sequence elements) and circularization yields according to circularization reaction (chemical PORA, enzymatic with T4 RNA ligase I, T4 RNA ligase II or autocatalytic PIE) with or without (w/o) splint DNA oligonucleotide.

**Figure 2.**
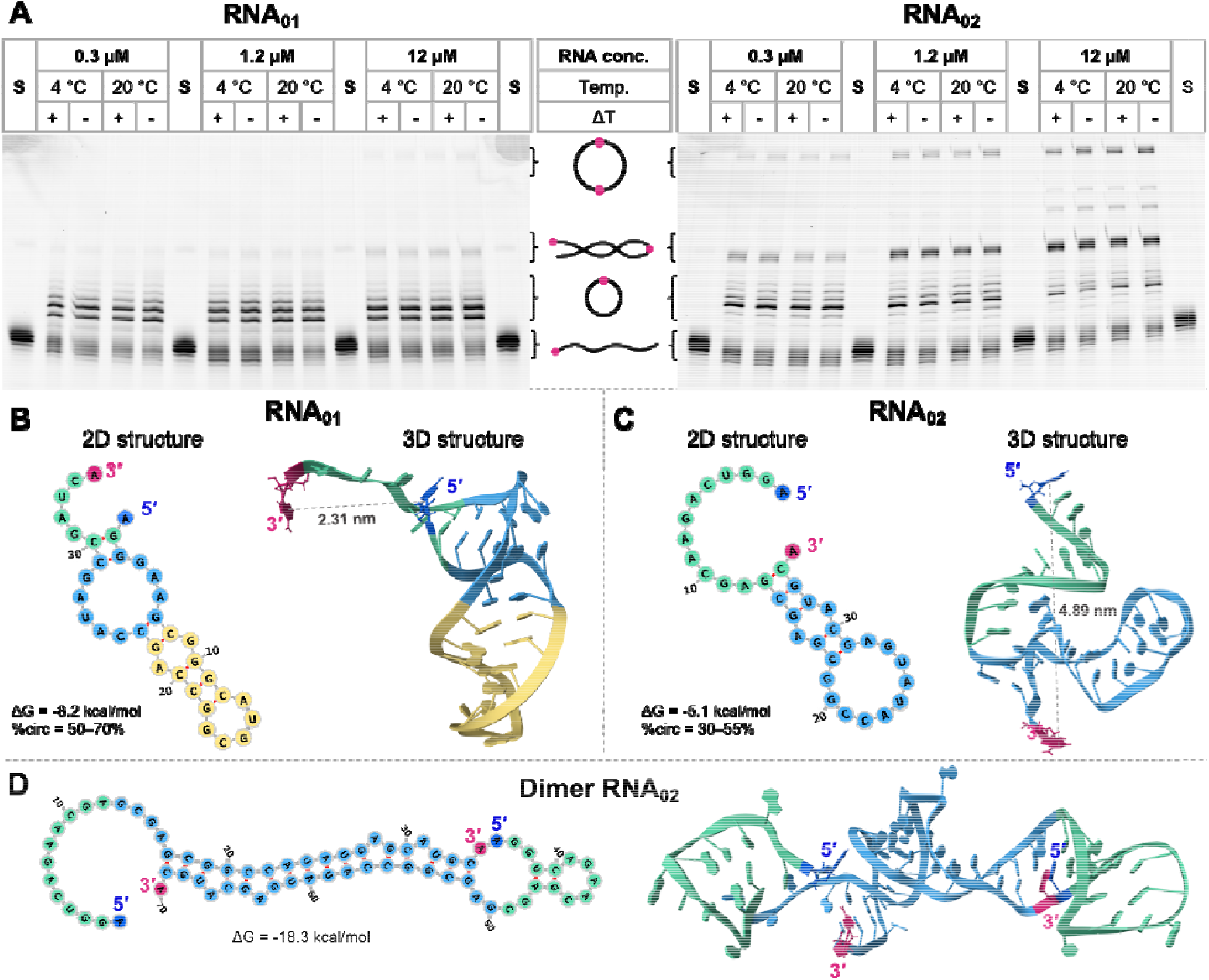
Optimization of chemical circularization of two RNA oligonucleotides (35 nt): A) PAGE analysis of the circularization reaction of RNA_01_ and RNA_02_. RNAs were circularized at concentrations of 0.3, 1.2, or 12 µM, at 4 or 20 °C, with (+ΔT) or without (-ΔT) thermal refolding (rapid heating and subsequent slow cooling) prior to circularization. S refers to an untreated RNA substrate. B-D) Minimum-free-energy (MFE) secondary structure (2D) models (predicted with RNAfold web server) and tertiary structure (3D) of predicted models (RNA Composer web server) for RNA_01_ (B), RNA_02_ (C), and RNA_02_ dimer (D).

To test the scope and limitations of this reaction, we first investigated the basic factors affecting chemical circularization. To that end, we tested the influence of pH, temperature, and RNA concentration on the circularization yield. We observed that the conversion of linear precursor (∼50%) was not significantly affected by pH (5.6–8.0) (Fig. S3), time, temperature (2 h at 20 °C or 16 h at 4 °C), or the concentration of the precursor (0.3–12 µM) (Fig. 2A). However, both RNA sequence and concentration were crucial for circularization selectivity. The circularization of E-RNA_01_ was, on average, more selective (50–70%) than that of E-RNA_02_ (30–55%), which also produced linear and circular dimers. Notably, the circularization of E-RNA_01_ did not lead to any linear and circular polymeric by-products (dimers and concatemers), especially at lower concentrations of RNA (Fig. 2A).

To gain insight into the sequence-dependence of RNAs circularization, secondary and tertiary structural models of the RNA_01_ and RNA_02_ sequences were generated using automated structure predictors (Fig. 2B-D).^27-29^ Computational predictions suggested that both RNA_01_ and RNA_02_ can form distinct secondary and tertiary structures. The structure of RNA_01_ features a stable hairpin (free energy of the thermodynamic ensemble -8.2 kcal/mol) with an open loop motif within which the 3□ and 5□ ends spatially converge. The predicted average distance between the 5□ and 3□ ends in the RNA_01_ model is shorter (2.3 nm), making the molecule prone to circularization (Fig. 2B). In contrast, the secondary structure of RNA_02_ formed a low stability hairpin (-5.1 kcal/mol). The 3□ and 5□ ends reside within a bigger open loop, with an increased end-to-end distance (4.9 nm). Moreover, the tertiary structure model of RNA_02_ suggested that the 3□ and 5□ ends could point in opposite directions (Fig. 2C). The spatial arrangement of these termini probably enhances the likelihood of intermolecular interactions rather than intramolecular ones, ultimately leading to the preferential formation of stable RNA dimers (-18.3 kcal/mol), especially at higher concentrations (Fig. 2A and 2D). Overall, the chemical circularization method worked well on the oligonucleotide models, and its selectivity and efficiency could be rationalized with the help of computational methods. Considering that the majority of long RNA sequences adopt tertiary structures with termini separated by no more than 10 nm, these results encouraged us to test circularization on longer RNAs.^30^

To that end, we performed chemical circularization of macromolecular RNAs (RNA_03_–RNA_08_, 276–760 nt), which included various protein-coding RNAs or their 5′ terminal fragments (Table 1). Here, we took advantage of a dinucleotide initiator with longer linker (EDA-PEG_5_-AG; Fig. S1). The efficiency of our chemical circularization for these RNAs was compared with enzymatic ligation of 5′-monophosphate RNAs catalyzed by T4 RNA ligases I and II. The chemical circularization process proceeded quite efficiently for most of the studied sequences, independent of their length. On average, chemical circularization (45–60%) was more efficient than the enzymatic ligation catalyzed by T4 RNA ligase I (5–40%; Table 1). We observed that the chemical circularization yield for RNAs longer than 900 nt was generally lower than for shorter RNAs (Table 1). Furthermore, the introduction of a polyA sequence (A_30_) at the 3′ end of the precursor resulted in a dramatic drop in circularization yield (<5%). For instance, the circularization efficiency of RNA_10_, which lacks the polyA (A_30_), was 33%, whereas for the same sequence with the polyA (RNA_11_), it was only 2%. A similar result was obtained for another sequence containing an A_30_ tract (RNA_09_) (Table 1, Fig. S4). The low efficiency of circularization for these sequences is likely due to the unstructured nature of the polyA region, which increases the 5′-3′ end-to-end distance in the precursor.^30^ Since our overall goal was to develop a circularization method applicable to all RNA sequences and lengths, we attempted to overcome this issue by applying a splint-mediated approach. In order to decrease the distances between the 5′- and 3′-terminal moieties in longer and unstructured RNAs, we annealed the linear precursors with oligonucleotide DNA splints (short DNA sequences that are complementary to the 5′ and 3′ flanking regions of the precursor RNA; Fig. 3). To that end, we applied computational modeling (RNAfold web server) to RNA_11_ (952 nt) containing human beta globin (HBB) 5′ and 3′ UTRs, *Gaussia* luciferase (GLuc) ORF, and A_30_ tract, to design four DNA sequences differing in length and targeting specific structural features of the precursor RNAs (ON1-ON4; 10-67 nt; Fig. 3A). Next, we synthesized a molecular probe consisting of RNA_11_ labeled with a fluorescent FRET pair, Cy5 and Cy3, at the 5□ and 3′ ends, respectively (Fig. 3B).^26^

**Figure 3.**
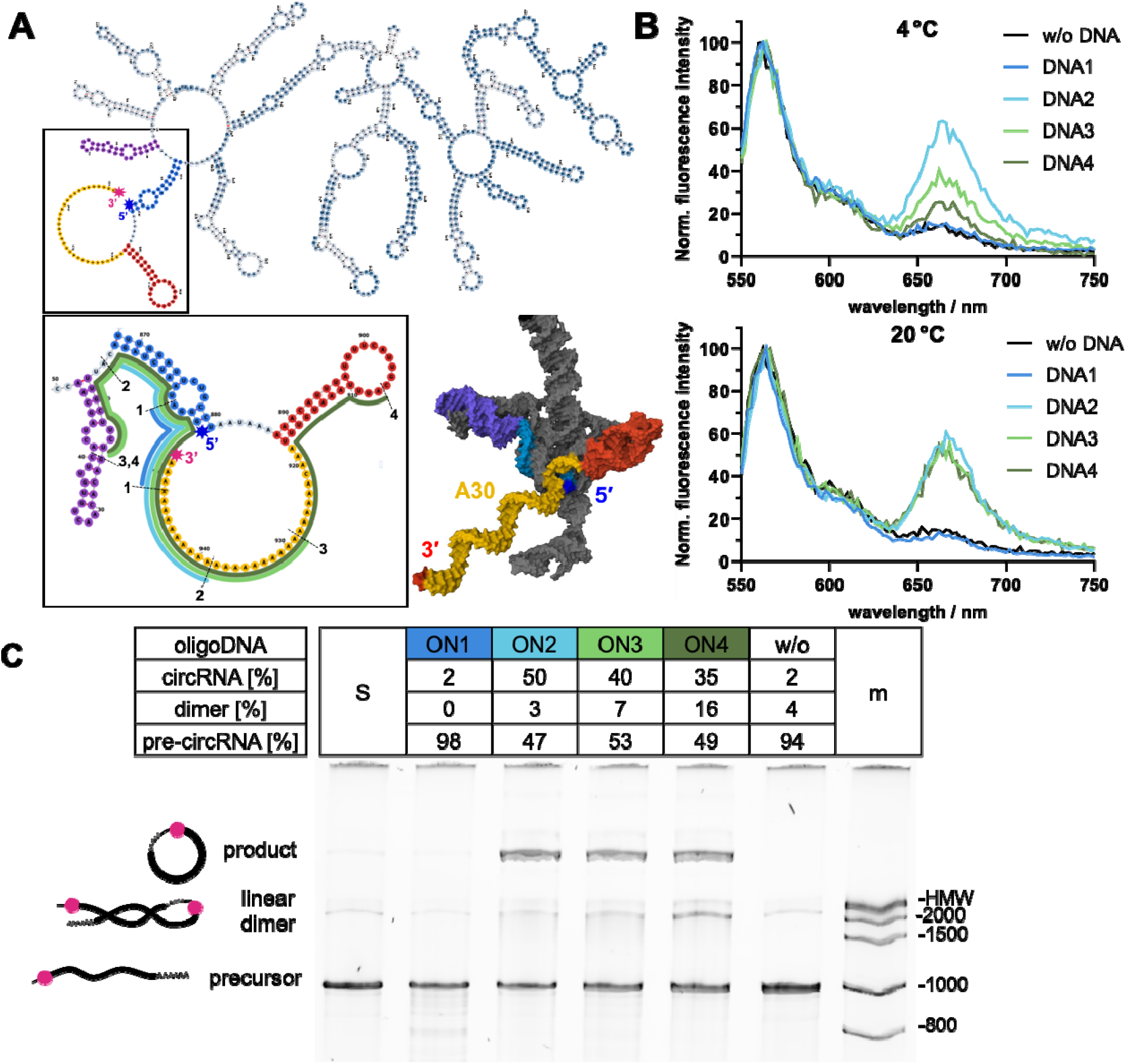
Circularization efficiency for challenging RNA sequences can be augmented using DNA splints: A) Optimization of splinted circularization, aided by the application of FRET probes and molecular modeling. The MFE secondary structure model of the RNA_11_ sequence and tertiary structure model of a selected fragment of the MFE secondary structure (predicted with RNAfold web server). Numbers 1–4 mark the regions of hypothetical interactions between RNA and oligonucleotide splints (ON1–4). B) Fluorescence spectra of a FRET probe (Cy5-RNA_11_-Cy3) after hybridization with an oligonucleotide splint (ON1–4) or without splint (w/o), recorded at 4 °C or 20 °C. C) PAGE analysis of the chemical product of RNA_11_ circularization, preformed after annealing the precursor with (ON1–4) or without (w/o) an oligonucleotide splint. S refers to untreated precursor.

Measuring and analyzing the fluorescence spectra recorded at different temperatures, before and after splint annealing with the probe, led to the selection of an optimal splint sequence (ON2, 28 nt; Fig. 3). For RNA_11_, the circularization efficiency in the presence of the optimal DNA splint (ON2, 28 nt, 50%) was over twenty-fold higher than that in the absence of DNA (2%) or in the presence of sub-optimal splints (2–40%, Fig. 3B,C). The use of longer DNA splints, ON3 (48 nt) and ON4 (67 nt), resulted in reduced circularization selectivity and increased linear dimer formation (Fig. 3C), whereas the shorter splint (ON1, 10 nt) did not improve circularization efficiency at all. The experiment revealed that the optimal splint has to form a sufficiently stable RNA-DNA duplex (melting temperature ∼50 °C) to unwind the less stable RNA-RNA secondary structure (melting temperature ∼20-30 °C), but should not be too long as it shifts the equilibrium toward dimer formation (Fig. 3A).

To verify the universality of this splint-based approach for longer RNAs, we applied it to six additional RNA sequences (RNA_09_, RNA_10_ RNA_12_-RNA_15_). Our observations indicated that the efficiency of chemical circularization, particularly for sequences possessing a polyA tract, increased significantly in the presence of a suitably designed splint.

Furthermore, during the circularization reaction, we observed little to no formation of polymeric side-products or nicking of the desired product. Overall, we found that splint-aided chemical circularization is equally efficient (44–63%) as the PIE methodology (34–54%, Fig. S5). In contrast, using T4 RNA ligase I for the circularization of long RNAs (>900 nt) resulted in little or no desired product (0–13%) and led to the formation of dimeric side-products. Circularization of the respective mRNAs with T4 RNA ligase II yielded higher circularization efficiencies (40–60%) than with T4 ligase I and were comparable to chemical circularization (Table 1). However, we noted that enzymatic ligation was significantly more sequence-dependent than chemical circularization.

To investigate how the quality of the linear precursor affects circularization efficiency, we conducted circularizations on RNAs purified by different methods. In vitro-transcribed (IVT) RNA containing an EDA motif [(EP_5_)RNA_13_, 1207 nt] was used as a model. After the IVT reaction, the crude mixture was purified using either a silica-based column (Monarch^®^ RNA Cleanup), a cellulose, or high-performance liquid chromatography (HPLC). The isolated linear (EP_5_)RNA_13_ was then subjected to a circularization reaction using the PORA protocol. The highest circularization efficiency was obtained for the sample purified by HPLC (55%) (Fig. S6A). RNA purified on cellulose resulted in a slightly lower circularization efficiency (46%) (Fig. S6A), while mRNA filtered through a silica column yielded the desired circRNA product with 49% efficiency (Fig. S6A). Overall, we established that the quality of the linear RNA precursor did not significantly affect circularization efficiency, which implies applicability of our protocol to in vitro-transcribed RNAs isolated using different routine protocols. It is known that dsRNA contaminants of in vitro-transcribed mRNA can trigger an undesired innate immune system activation and strongly affect the outcomes of biological evaluations.^31^ Therefore, we also tested the obtained linear and circular RNAs for the presence of dsRNA (Fig. S6B). As expected, both HPLC and cellulose purification efficiently removed dsRNAs for both linear and circular samples, whereas the purification on silica-based column did not.^32-34^ Hence, dsRNA contaminants in chem-circRNAs can be removed using solutions previously developed for linear RNAs, making our method scalable and suitable for existing manufacturing pipeline.

### Analytical methods for identifying and isolating chem-circRNA

The purity and homogeneity of circular RNA are crucial for mitigating an innate immune response and ensuring sustained protein production. Depending on the circularization method used, the resulting mixture may contain remnants of unreacted linear precursor, polymerized linear and circular side products such as dimers and concatemers, excised intron fragments, nicked circRNA, or splint templates that can activate the innate immune response.^32^ To establish optimal methods for identifying and purifying chem-cricRNA, we first reviewed established methods from the literature. Then, we conducted a detailed examination and optimization of selected methods, leading to significant improvements and insightful findings.

The yields of RNA circularization and the homogeneity of circRNAs were determined based on electrophoretic analyses. Initially, we attempted to resolve linear and circRNAs on denaturing agarose gel. However, we typically observed poor resolution or overlap between linear and circular species of corresponding weight, which has also been reported previously.^5,6,14,37^ Consequently, we turned our attention to denaturing polyacrylamide gel electrophoresis (PAGE). By varying both the degree of cross-linking of the polyacrylamide (PAA) and the final concentration of the AA/MBAA blend, we determined the optimal density and cross-linking degree of polyacrylamide gels for the best migration and separation of linear and circRNA (Fig. 4). In all studied cases, the migration speed of the circular RNA macromolecules in polyacrylamide gel was significantly slower than that of linear precursors of corresponding weight (Fig. 4A) or high-molecular-weight (HMW, 3000–6000 nt) linear RNA molecules (Fig. 4B, C). A higher gel density (20% PAA) with a cross-linking degree of 5% (19:1 AA/MBAA) favored the separation of short linear mRNAs and their circularization products (RNA_01_, 35 nt) (Fig. 4A). A lower gel percentage, corresponding to an increase in pore size, enhanced the migration rate of nucleic acids, as demonstrated for RNA_03_ (2 6 nt) and RNA_04_ (760 nt) on a 15% and 6% PAA gel, respectively. Conversely, a higher degree of cross-linking (19:1 AA/MBAA) significantly inhibited the migration of circRNA_03_ (276 nt) and circRNA_04_ (760 nt). Reducing the degree of cross-linking to 2.3% (38:1 AA/MBAA) increased the migration rate of circRNA while maintaining substantial separation between linear and circular RNA topologies. Such pronounced differences in nucleic acid migration are primarily attributed to variations in secondary structure rather than molecular weight. Through these experiments, we demonstrated that a 6% PAA with a 38:1 AA/MBAA ratio provided the best separation of linear and circular RNA species for most RNAs. Using these conditions, we were able to access the composition of the reaction mixtures in a simple, precise, and reproducible manner.

**Figure 4.**
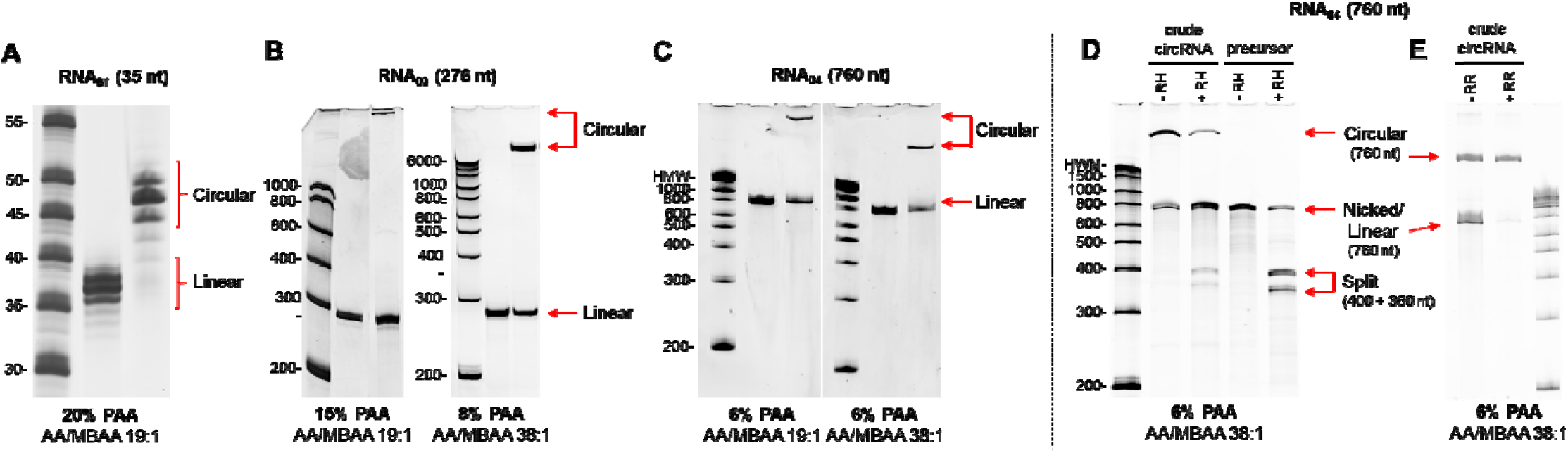
Electrophoretic and enzymatic methods for the analysis of chem-circRNA: A-C) Resolution of circular RNAs and their linear precursors based on molecular size (A: RNA_01_ 35 nt, B: RNA_03_ 276 nt, C: RNA_04_ 760 nt), gel density, and crosslinking degree of denaturing polyacrylamide. D, E) Treatment of crude reaction mixtures with endo- and exonucleases (RNase H - RH and RNase R – RR, respectively) to confirm the circular topology of the reaction products.

On the other hand, the topology of circRNAs can be readily verified by treating them with endonucleases and exonucleases, including RNase H and RNase R.^3,38,39^ Digestion of intact chem-circRNA with oligonucleotide-guided RNase H predominantly produced a single band, in contrast to the nicking of its linear precursor, which yielded two split products of lower molecular weight (Fig. 4D). RNase R, with 3′ to 5′ exonuclease activity, eliminates linear contaminants from circRNA preparations.^3,4^ Due to the unique close-loop structure, circRNAs remain resistant to RNase R digestion, which has been used for circRNAs enrichment. We found that chem-circRNAs are essentially also resistant to RNase R, as the RNase-treated circularization reaction yielded solely the chem-circRNA band (Fig. 4E). However, when applying RNase R digestion for chem-circRNAs enrichment, we observed that both the quality of the circRNA sample and the presence of some sequence and structural elements may affect the selectivity of the reaction (Fig. S7). Digestion of RP-HPLC-purified mixtures of circular and linear RNA significantly increased the selectivity of RNase R towards the linear RNA species (Fig. S7A). To investigate the resistance of chemically circularized RNAs to RNase R, we conducted RNase R digestion on three chem-circRNA_11_ variants with different linker lengths [(EDA-RNA_11_, EP_5_-RNA_11_ and EP_5_Cap1-RNA_11_], as well as an enz-circRNA_11_ [(p)RNA_11_] with a phosphodiester bond at the junction site. We established that the introduction of a chemical linker in chem-circRNA_11_ does not preclude the use of RNase R for chem-circRNA enrichment (Fig. S7B). However, we also observed undesired degradation of circRNAs_11_. This issue may be resolved by optimizing RNase R digestion conditions, such as enzyme concentration incubation times or buffer composition.

To obtain high-purity circRNA samples, we employed PAGE separation of linear and circular RNA. The chem-circRNA molecules were resolved on PAA gel and extracted using either the crush-and-soak method or electroelution. We found that the UV spectra of the isolated material differed greatly, indicating that crush-and-soak isolation leads to contamination of RNA with PAA. In contrast, electroelution yielded chem-circRNA of high purity (>90% on PAGE) with an acceptable UV spectrum (Fig. S8A, B). Given that RNAs separated on denaturing agarose gels are reportedly non-translatable due to interactions between denaturants (formaldehyde) and nucleotide monomers,^40^ we investigated the translational activity of circRNAs isolated from polyacrylamide gels (Fig. S8C). The circ(EP_5_)RNA_13_ and reference circular RNA obtained by PIE splicing strategy (circRNA_16_-PIE) were purified via PAGE/electroelution or RNase R treatment to compare their translational activity in cultured cells. The biological impact of circRNA isolation was assessed in three cell lines (A549, HEK293T, HepG2). Regardless of the circRNA sample, total protein levels were comparable between the isolation methods (Fig. S8C).

Another possible method for chem-circRNA purification is isolation from the reaction mixture by high-performance iquid chromatography (HPLC). We investigated this in three modes: ion-exchange chromatography (IEC), size-exclusion chromatography (SEC), and reversed-phase chromatography (RP) (Fig. 5A). Previous studies utilized SEC for isolating circularization products from self-catalytic precursors using permuted intron-exon (PIE) methodology.^5,6,14^ SEC can be effective in such scenario because during the transesterification reaction, the linear precursor eliminates chain fragments corresponding to sequences of autocatalytic introns, contributing to a decrease in molecular mass. However, in our hands, SEC did not facilitate the separation of linear and circular RNA molecules of the same size and sequence (such as chem-circRNA and its precursor or the desired circRNA and nicked product). Successful separation of RNA molecules based on their topology was achieved only in RP mode in the presence of ion-pairing agents (IP). Optimal separations were achieved at 60 °C temperature on polystyrene divinylbenzene copolymer resin in the presence of n-hexylammonium acetate and acetonitrile (Fig. 5B). Notably, the IP-RP-HPLC method is effective only for RNAs shorter than 1000 nt; for longer RNAs, the retention times for circular and linear forms are similar, hindering efficient separation. The combination of IP-RP-HPLC and PAGE provided us with a set of tools to precisely analyze circularization reactions and isolate the desired circRNA products with high purity (Fig. 5C) .

**Figure 5.**
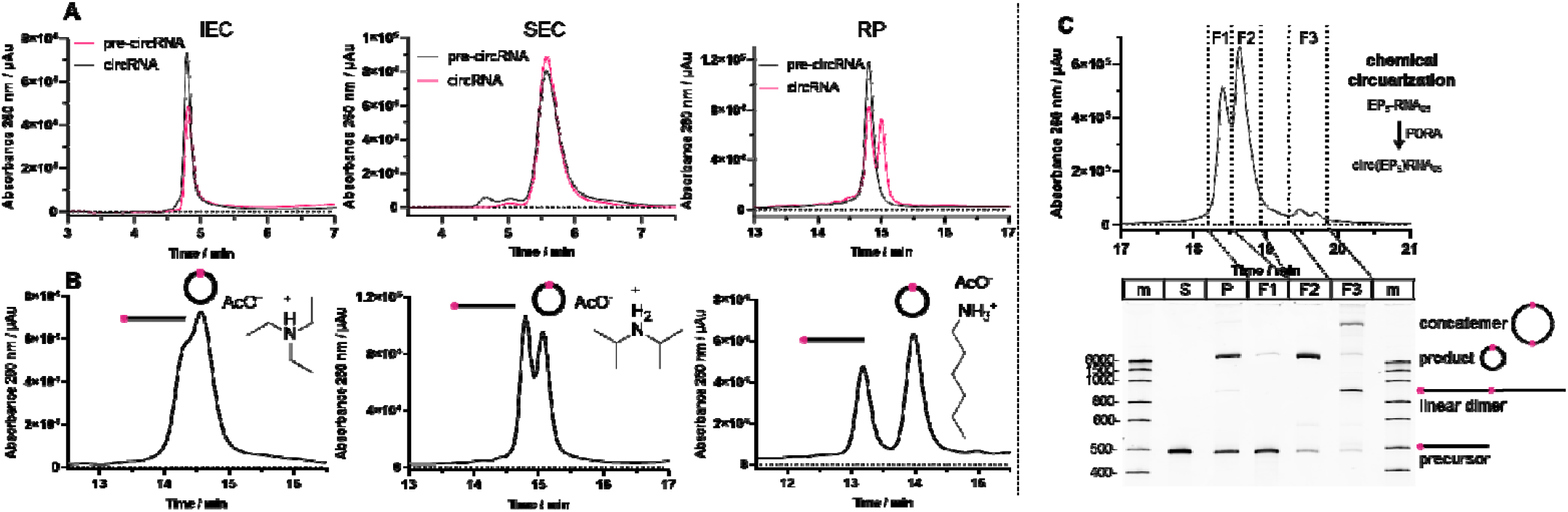
Analysis and purification of chem-circRNA can be achieved using high-performance liquid chromatography (HPLC): A) Samples of (p)RNA_04_ (pre-circRNA_04_) and crude products of its circularization [circ(p)RNA] were resolved using different HPLC methods: size-exclusion (SEC), ion-exchange (IE), and reversed-phase (RP). B) RP-HPLC separation of linear (lower retention time) and circular (higher retention time) RNA_06_ (598 nt) sequence in dependence of the ion-pairing reagent used (triethylammonium acetate, diisopropylammonium acetate, or hexylammonium acetate). C) Separation and analysis of the circularization products (chemical for RNA_05_) with RP-HPLC and PAGE. Fractions labeled as F1 contains mostly precursor RNA, F2 comprises the circular product of high purity, and F3 includes linear dimers and concatemers.

### Translation of chemically modified circRNA

After establishing a robust methodology for preparing of chem-circRNAs, we applied this technique to investigate the influence of RNA sequence and chemical modifications on the biological activity of mRNA. Our research focused on two main areas: (1) the biological activity and stability of chemically circularized mRNA, and (2) the incorporation of the (m^7^G) cap structure into chemically circularized mRNAs and its biological impact. To explore the translation of chem-circRNA analogs and their structure-activity relationship, we synthetized a series of precursor sequences (RNA_09_, RNA_12_–RNA_15_), encoding a *Gaussia* luciferase, flanked by 5D and 3′ UTRs and polyA sequences (Table 1), which were chemically circularized. The resulting chem-circRNA analogs were equipped with either a PEG_5_ linker [circ(EP_5_)RNA] or an endocyclic m^7^G cap [circ(EP_5_Cap)RNA]. All chem-circRNAs and their linear precursors were transfected into four different cell lines (A549, HEK 293T, HepG2, and HeLa).

First, we investigated whether the introduction of the (EP_5_)EDA chemical linker impaired the functionality of circRNAs in any way (Fig. 6A-D). We used model mRNAs incorporating the EMCV IRES sequence, which facilitates cap-independent translation initiation, to test this. To avoid bias from the translation of linear RNA species (circRNA precursors and nicked RNA circles), we prepared circRNA sequences without 3□ UTR or structure-stabilizing elements (circRNA_12_) (Fig. 6A, B). This approach ensured that the translational activity of linear counterparts was negligible. The activity of these RNA constructs was compared to unmodified circRNAs obtained by enzymatic circularization of the same sequence (Fig. 6A, B). Additionally, circRNA obtained by PIE (circRNA_16_-PIE) and linear mRNA optimized for cap-dependent translation (Cap1-RNA_11_) served as references (Fig. 6B). As expected, the translational activity of circRNAs was much higher than that of their linear precursors (Fig. 6B and S9). Importantly, chemically obtained circRNA showed comparable translational activity and stability to enzymatically obtained circRNA (Fig. 6A, B).

**Figure 6.**
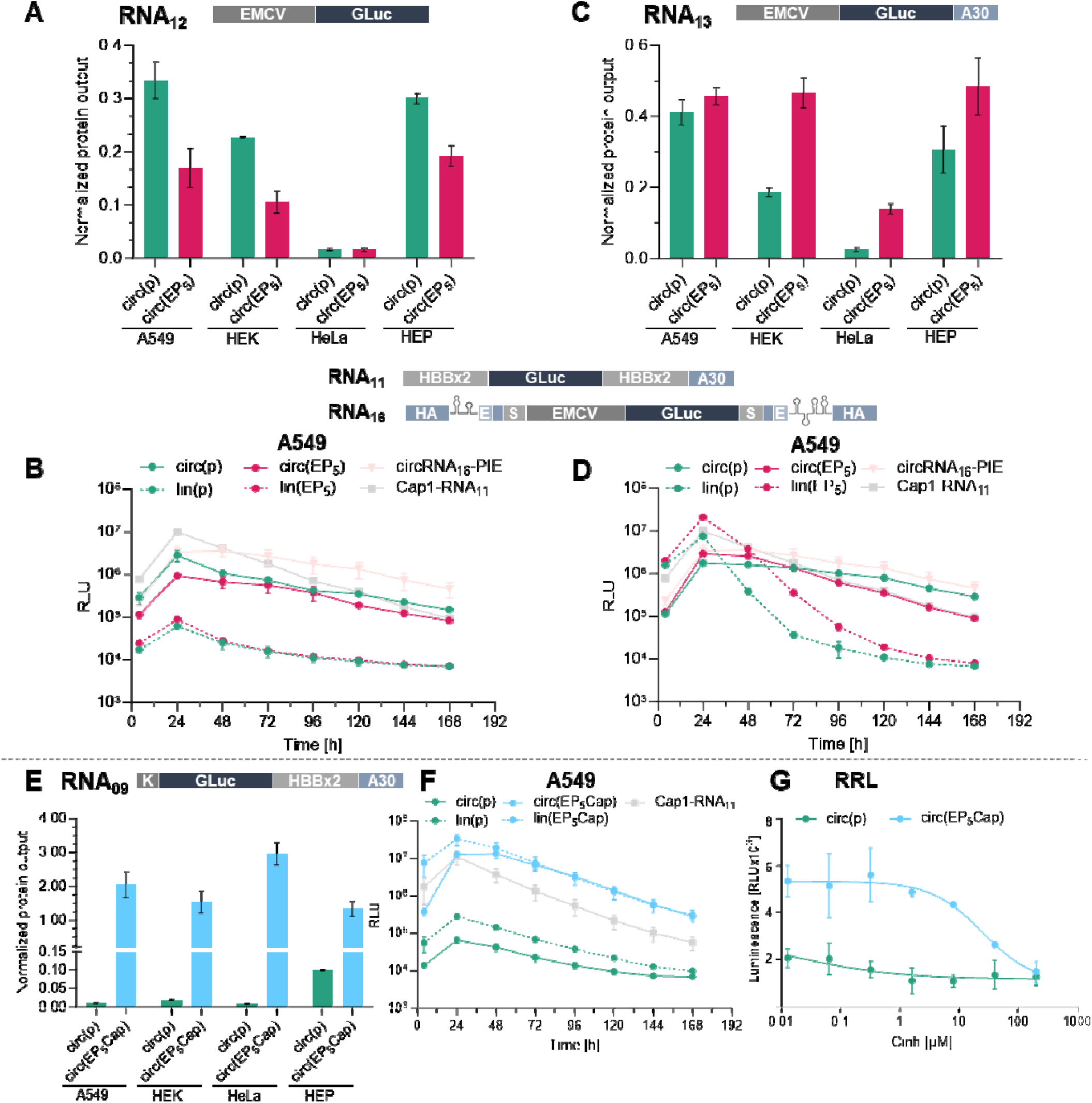
Chem-circRNAs can undergo translation in living cells via IRES and cap-dependent mechanisms: A-F) Protein expression profiles of linear and circular mRNAs of different sequences: RNA_12_ (A, B), RNA_13_ (C, D), RNA_09_ (E, F). Cells of four different lines [A549, HEK 293T (HEK), HepG2 (HEP), and HeLa] were transfected with circRNA coding *Gaussia* luciferase derived from linear precursors bearing 5′ monophosphate (p-) or EDA-PEG_5_Cap (EP_5_Cap-) structure, or their precursors. The bars represent the relative evels of the reporter protein in cell culture medium, determined at the indicated time points and normalized to the level obtained for the reference canonical mRNA (Cap1-RNA_11_). Data represents the mean values and standard deviation, n=3. G) Translation of circ(EP_5_Cap)RNA_09_ in rabbit reticulocyte lysate (RRL) is inhibited in the presence of the cap-dependent translation inhibitor (m^7^Gp_s_ppG), confirming the cap-dependent translation mechanism.^41^

Next, we investigated the impact of elements required for cap-dependent translation, such as the polyA tract, on the durability of protein production and the translational activity of our chem-circRNA analogs. The effect of polyA (A_30_) was studied using circRNA_13_ (Fig. 6C, D). As expected, the introduction of the polyA tract led to enhanced stability of the linear precursors, resulting in a burst of protein expression within the first 24 hours post-transfection, followed by a rapid decline in protein levels (Fig. 6D and S9). In contrast, both circ(EP_5_)RNA_13_ and circ(p)RNA_13_ provided sustained protein production, similar to circRNAs without the polyA (circRNA_12_) (Fig. 6D, B). Moreover, the duration of expression in cells for all circRNA analogs (expression halftime 29–58 h) was increased compared to their linear precursors (8–11 h) (Fig. S10). Overall, the data suggest that, unlike linear RNA, the presence of a polyA tract has a negligible influence on circRNA durability. We also observed that both chemically and enzymatically circularized circRNAs exhibit a protein production profile similar to that of circRNA_16_-PIE, which contains a custom-designed spacer at the splicing site.^5, 6^

Next, we used our circularization method to explore the implications of the m^7^G cap in the context of circRNA translation (Fig. 6E, F). To promote cap-dependent translation of circRNA, we circularized mRNA containing an EDA-PEG_5_-m^7^GpppA_m_G cap structure (EP_5_Cap), a minimal 5□ UTR (Kozak), 3□ UTR (from human beta-globin gene, HBB), and a 30 nt polyA tract [circ(EP_5_Cap)RNA_09_]. The cap functionality in chemically circularized mRNA [circ(EP_5_Cap)RNA_09_] was compared with translationally inactive circ(p)RNA_09_ produced enzymatically (Fig. 6E and F). As before, linear Cap1-RNA_11_ served as a reference. The introduction of the cap structure into the circRNA molecule dramatically increased the protein levels (200-fold on average) compared to the enz-circRNA (Fig. 6E, F). Surprisingly, the discrepancy in translation efficiency between the capped circRNA and its linear precursor was relatively small (2-fold lower on average, Fig. 6F and S9). Furthermore, both linear and circular capped RNAs displayed similar protein production profiles over time (Fig. 6F). To confirm that the capped circRNA analogs translate in a cap-dependent manner, we compared the expression of circ(EP_5_Cap)RNA_09_ and circ(p)RNA_09_ in a cell-free system (rabbit reticulocyte lysate, RRL) in the presence of a cap-dependent translation inhibitor (m^7^Gp_s_ppG dinucleotide, Fig. 6G).^41^ Indeed, increasing the inhibitor concentration decreased the level of reporter protein from the translation of circ(EP_5_Cap)RNA_09_, but not of circ(p)RNA_09_. This strongly suggests that the endocyclic m^7^G cap structure in this chem-circRNA construct is fully functional, although the durability and translatability of the circular and linear capped mRNA species are very similar.

Then, we investigated the coexistence of both m^7^G cap structure and IRES in the context of circRNA translation. We hypothesized that the interplay between these distinct mechanisms of translation initiation in circRNAs might enhance overall protein production. To test this, we circularized mRNAs containing a 5□ EDA-PEG_5_Cap, followed by EMCV IRES and GLuc reporter sequences alone (RNA_12_) or with different downstream elements such as a polyA tract (RNA_13_), HBB 3□ UTR (RNA_14_), or HBB 3□ UTR followed by a polyA tract (RNA_15_) (Fig. 7A-D and S9). Linear Cap1RNA_11_ served as a reference.

**Figure 7.**
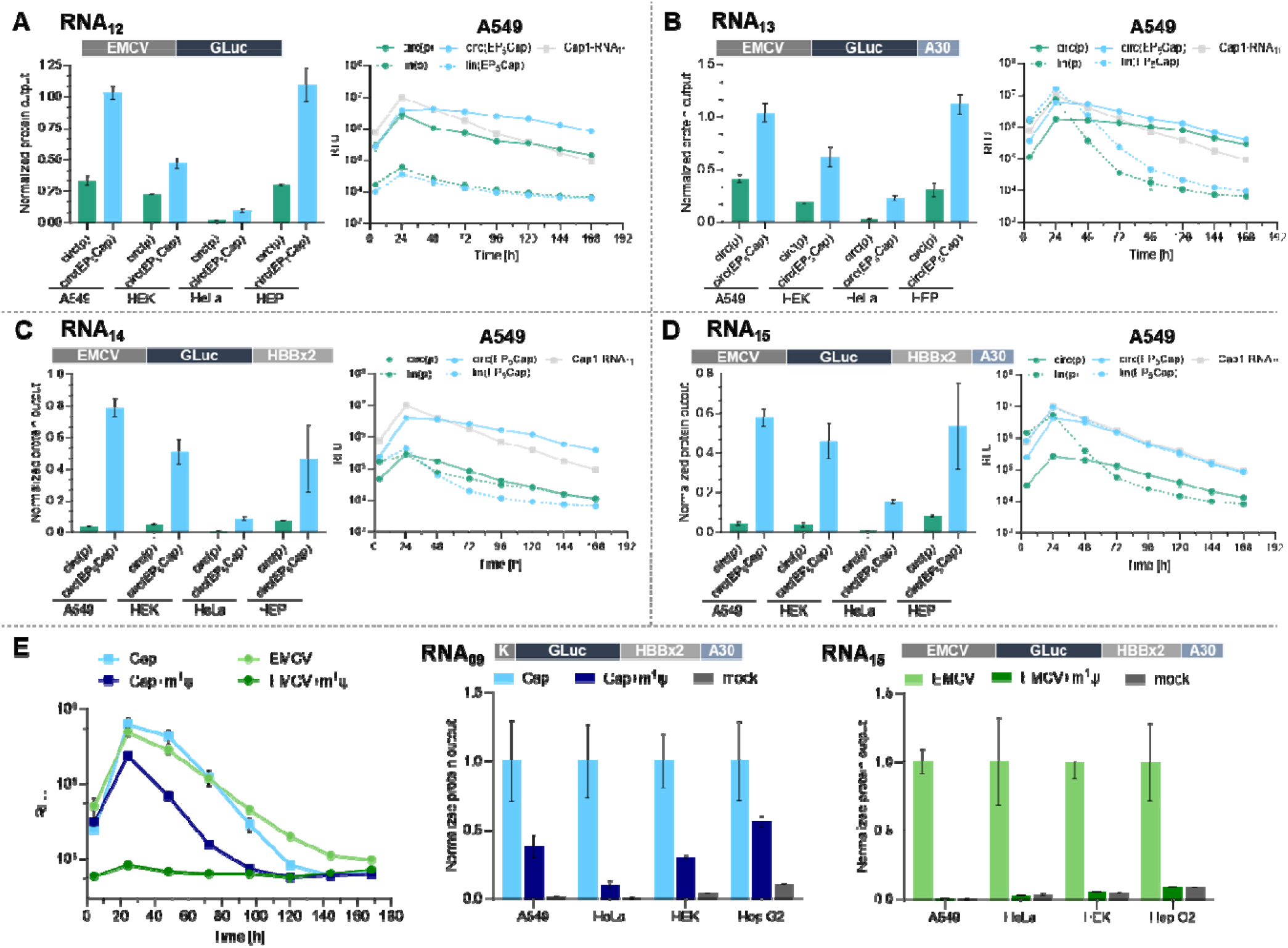
Protein expression profile of linear and circular mRNA analogues: A) RNA_12_, B) RNA_13_, C) RNA_14_, D) RNA_15_. Cells of four different lines [A549, HEK 293T (HEK), Hep G2 (HEP), and HeLa] were transfected with circRNA derived from linear precursors bearing 5′ monophosphate (p-) or EDA-PEG_5_Cap (EP_5_Cap-) structure and coding *Gaussia* luciferase (RNA_12_ – RNA_15_ sequences). The bars represent the relative levels of the reporter protein harvested over 168 hours of cell culture, normalized to the level obtained for canonical mRNA reference (Cap1-RNA_11_). Data represents the mean and standard deviation, n=3. E) Impact of N1-methylpseudouridine introduction on the translation of circRNA. Cells were transfected with circ(EP_5_Cap)RNA_05_ (Cap), circRNA_15_ (EMCV), and their counterparts containing N1-methylpeudouridine (+m^1^Ψ). The bars represent relative levels of the reporter protein harvested over 168 hours of cell culture, normalized to the level obtained for non-modified Cap- and IRES-containing circRNA reference [circ(EP_5_Cap1)RNA_15_ circRNA_15_]. Data represents the mean values and standard deviation, n=3.

The integration of both the cap and IRES resulted in a modest improvement in protein output compared to enzymatically circularized RNAs, both with and without a polyA downstream of the reporter gene (3-fold and 5-fold on average for sequences RNA_12_ and RNA_13_, respectively, Fig. 7A, B). Surprisingly, the extension of the 3□ UTR region resulted in an even more prominent surge in the protein output of capped circRNAs, with an average increase of 13-fold and 12-fold for sequences RNA_14_ and RNA_15_, respectively (Fig. 7C, D). Importantly, the introduction of the endocyclic cap structure in chem-circRNA did not disrupt the trend of extended protein production, unlike its linear counterpart (Fig. 7C, D). Taken together, these results indicate a supportive, though nuanced, influence of the endocyclic cap structure on translation of EMCV-IRES-containing circRNA.

N1-methylpseudouridine (m^1^Ψ) is among the most critical modifications in therapeutic mRNA. It has been shown to decrease the activation of cytosolic and toll-like receptors (TLRs), thereby lowering cellular immune response and mRNA toxicity.^42^ On the other hand, the introduction of m^1^Ψ and other noncanonical nucleobases can disrupt the function of RNA elements, such as IRES or autocatalytic introns, by altering their secondary structure.^6^ We decided to investigate whether an endocyclic cap structure could replace the IRES element and facilitate the translation of circRNA containing the m^1^Ψ base modification. For this, we prepared linear precursors by IVT in the presence of m^1^Ψ-triphosphate and performed either chemical or enzymatic circularization. The circRNAs were then transfected into four cell lines (Fig. 7E). As expected, IRES-dependent translation of circRNA (EMCV+m^1^Ψ) obtained by enzymatic circularization was abolished. In contrast, the chemically obtained capped circRNA (Cap+m^1^Ψ) with m^1^Ψ and HBB UTRs remained translationally active, albeit its activity was decreased compared to the chem-circRNA without m^1^Ψ. Overall, these results demonstrate that the developed chemical circularization is compatible with RNA base-modifications and opens new avenues for optimizing the biological activity of circRNA.

## DISCUSSION

In this work, we report the first method for the chemical circularization of in vitro transcribed (IVT) mRNA. Exogenously delivered circRNA represents a promising drug modality due to its extended lifespan in the cytosol. Since circRNAs lack the free ends, they are not susceptible to exonuclease-mediated degradation, which is the main pathway for the bulk removal of cytoplasmic RNA.^2,43^ To date, the most well-characterized methods for circularizing linear mRNA involve enzymatic ligation (RNA or DNA ligases) and ribozymatic methods based on self-splicing introns (Fig. 1).^5,8-14,5,15^ Chemical methods have been limited to short RNA sequences applicable to rolling circle amplification (RCA) of short peptides.^44^ Our group has been involved for many years in developing chemical modifications of RNA ends. In this work, we leveraged this experience to develop a method of joining modified 5′ and 3′ ends of in vitro transcribed RNA, i.e. to chemically circularize full-length mRNAs. We were inspired by recent observations, made by us and others,^26,30^ that even in long RNA macromolecules, the 5′ and 3′-ends are usually sufficiently close to each other to enable fluorescence energy transfer (<10 nm), which should mean they are also close enough to efficiently undergo an intramolecular chemical reaction.

As a result, our study describes a novel method for obtaining chemically circularized RNAs (chem-circRNAs) based on a direct and selective chemical reaction called PORA (Periodate Oxidation and Reductive Amination). The starting material for this reaction are RNAs obtained by standard in vitro transcription in the presence of transcription primers carrying an ethylenediamine motif. PORA is performed on the RNA product in a one-pot two step reaction relying on simple reagents, taking less than 2 h and providing conversions in the range of 35-63%. Notably, our methodology allows the implementation of chemical modifications into circRNAs that were previously inaccessible to this modality, as demonstrated by incorporating an endocyclic cap structure capable of driving cap-dependent translation and m^1^Ψ into circRNA. Since typical circRNAs lack the m^7^G cap, their design has been limited to the presence of certain sequences that induce internal translation initiation, such an internal ribosome entry sites (IRES) or short IRES-like A/U enriched sequences^35, 36, 45^ Our chemical approach thus opens completely new possibilities for the design, modification and fine-tuning of circRNAs.

Using simple 35 nt RNA models (RNA_01_ and RNA_02_), we demonstrated that circularization yields are predominantly dependent on the local secondary structure of RNA (end-to-end distance) rather than its length (Fig. 2). We further showed that circularization is equally efficient for RNA of various lengths and sequences, up to almost 1500 nt long. A reduction in circularization efficiency was observed for longer RNAs and those with unstructured 3′ ends (incorporating 3′ terminal polyA tracts). This issue was addressed by using secondary structure control with oligonucleotide DNA splints, achieving circularization efficiencies ranging from 40% to 60%, even for complex macromolecular RNA constructs (Table 1). We also developed methods that enable efficient separation of chem-circRNAs from their unreacted linear precursors and provided a set of molecular tools to verify the structure and homogeneity of chem-circRNAs. Our results demonstrate that chem-circRNAs undergo translation in living cells similarly to enzymatically obtained circRNAs. Moreover, the duration of expression in cells for all chem-circRNA analogs was extended compared to their linear precursors (Fig. S10). We also investigated the impact of elements required for cap-dependent translation, such as the m^7^G cap and the polyA tract, on the durability of protein production and the translational activity of our chem-circRNA analogs. The addition of polyA (A_30_) tract did not significantly affect the biological properties of circRNAs (Fig. 6A-D), whereas the addition of the m^7^G cap increased translational activity, although the effect was sequence dependent. We demonstrated that the endocyclic cap in chem-circRNA maintains its functionality, as evidenced by enhanced protein production compared to unmodified circRNAs (Fig. 6E and 6F) and susceptibility to inhibition by compounds targeting the cap-binding translation factor, eIF4E. Overall, our data reveal that chemical circularization is compatible with RNA base-modifications and that the endocyclic cap moiety could serve as an alternative to IRES for promoting circRNA translation (Fig. 7E). However, further extensive optimization of chem-circRNA sequences and chemical structures is crucial to fully realize the potential of this methodology. Nonetheless, to our best knowledge, this work describes several unprecedented achievements in RNA circularization: (i) the chemical circularization of mRNA encoding full-length protein, (ii) the chemical circularization of IVT RNA, (iii) the development of circular RNAs that undergo cap-dependent translation, and (iv) the circularization of m^1^Ψ-modified RNA. In contrast to enzymatic and autocatalytic circularization, our approach allows for site-selective introduction of RNA chain modifications simply by adding an analog of a substrate for RNA polymerase during the IVT reaction. This gives users a freedom of choice of chemical modifications that could help expand the repertoire of known and tested circRNA modifications. In light of that, we believe this work opens an unexplored avenue for capped circular RNA and potentially a new generation of RNA therapeutics.

## METHODS

More details on experimental procedures can be found in the Supplementary Data.

### Preparation of linear 5′ end modified and unmodified mRNA as a circularization precursors

The synthesis of the respective IVT initiators and intermediates is detailed in the Supplementary Data.

Unmodified and modified linear mRNA precursors were synthesized by in vitro transcription from a linearized plasmid DNA template using T7 RNA Polymerase, HC (Thermo) followed by treatment with DNAse I (Thermo). mRNA was purified using a silica-based column followed by RP-HPLC.

### Chemical circularization

An aqueous solution of the precursor RNA (30-40 µg in 490 µl, ∼140 µM) was diluted with buffer (70 µl, 800 mM KH_2_PO_4_, 200 mM NaCl, pH 7.0) and incubated at 65 °C for 5 min, followed by cooling to r.t.. After 10-15 min at r.t., a fresh solution of sodium periodate (70 µl, 10 mM) was added, and the reaction mixture was immediately placed in dark and incubated at 25 °C for 30 min. A fresh solution of sodium cyanoborohydride (70 µl, 200 mM) was added, and the incubation was continued for 90 min. The crude circRNA product was on a silica based-column (Monarch RNA cleanup kit, 50 µg, NEB) followed by RP-HPLC. For additional purification and removal of linear RNA, the product was treated with RNase R as described below (see section RNase R digestion).

### Chemical circularization with complementary splint

The reaction was carried out following the procedure outlined above, with the modification that a complementary oligonucleotide (∼11 µl, 10 µM, 1.5 eq, see Supplementary Table 2 for sequence) was employed.

### RNase R digestion

A solution of RNA (43 µl, 10 µg) was incubated at 65 °C for 5 min and rapidly cooled in ice bath. After 5 min at 0-4 °C, RNase R buffer ×10 (5 µl, 200 mM Tris-HCl, 1 M KCl, 1 mM MgCl_2_, pH 7.5, ABM), RiboLock RNase inhibitor (1.25 µl, 40 U/µl, Thermo), and RNase R (1 µl, 10 U/µl, ABM) were added. After 30 min of incubation at 37 °C the products of digestion were analyzed with PAGE directly or isolated from the reaction mixture using Monarch RNA cleanup kit (10 µg, NEB).

### RNase H nicking analysis

Circularization reaction crude after HPLC purification was heated at 65 °C for 5 min in the presence of a DNA probe (Supplementary Data) and gradually cooled to 37 °C for 30 min. Next, RiboLock RNase inhibitor (0.5 µl, 40 U/µl, Thermo), and RNase H (1 µl, 5 U/µl, Thermo) were added. After 30 min of incubation at 37 °C the products of digestion were directly analyzed with PAGE.

### Translation inhibition in RRL system

Flexi Rabbit Reticulocyte Lysate (4 μL, Promega) was diluted with 4 μL solution containing amino acid mixture (100 μM), KOAc (500 mM, and MgCl_2_ (2.5 mM), and incubated for 1 h at 30 °C. Next, 2 μL of RNA (50 ng/µl) mixed with m^7^GpsppG (0–200 μM) was added to 8 μL of the lysate mixture and incubated for 1 h at 30 °C. Next, the reaction was immediately placed into a microplate reader (Synergy H1, BioTek), 50 μL of h-coelenterazine in 1× PBS was added and the luminescence was measured at 25 °C. The assay was performed in duplicate to calculate the and mean and standard error (SEM).

### Cell culture and transfection (A549, HEK 293T, HeLa, Hep G2)

One day before transfection, 8×10^3^ cells were seeded per well of a 96-well plate and incubated under standard conditions (37 °C, 5% CO_2_, saturating humidity). Transfection was performed with Lipofectamine Messenger MAX (Invitrogen, Waltham, MA, USA), according to the manufacturer’s instructions. Each RNA variant (65 ng) was transfected in triplicate. At appropriate time points, 20 µL of medium from each well was withdrawn to test the amount of produced protein, and the remaining medium was completely replaced with fresh one. The amount of luciferase released into the medium was analyzed using the GLuc GLOW Assay (NanoLight Technologies, Norman, OK, USA) reagents, according to the manufacturer’s instructions using an EnVision plate reader (Perkin Elmer, Waltham, MA, USA)

## Supporting Information

Supplementary figures and tables, full DNA sequences, detailed descriptions of chemical syntheses, characterization of chemical compounds, experimental procedures involving *in vitro* transcription, RNA circularization and preparation, nucleolytic assays, FRET probes, cell culture and cell-free experiment are available.

## AUTHOR INFORMATION

## AUTHOR CONTRIBUTION

**JJ, JK** Conceived the research and acquired funding for this project, Supervision, Writing - Review & Editing. **AM** and **MWK** contributed equally. **JJ, JK, AM** Conceptualization, **AM, MWK** Methodology, Validation, Investigation, Visualization, Resources **AM, MWK, JK** Writing - Original Draft. **KC** Investigation, Validation. **KF** Investigation, Visualization, Resources. **TS** Investigation, Resources, **MW** Resources. **DN, JG** Provided useful comments and feedback for the manuscript, Supervision. All authors have given approval to the final version of the manuscript.

## ACKNOWLEDGMENT

The authors are grateful to Katarzyna Prokop and Andrzej Dziembowski (International Institute of Molecular and Cell Biology in Warsaw) for their support in plasmids preparation and helpful discussions. This work was supported by the Virtual Research Institute “HERO” project from Polish Science Fund.

## CONFLICTS OF INTERESTS

The authors declare no competing financial interest.

---

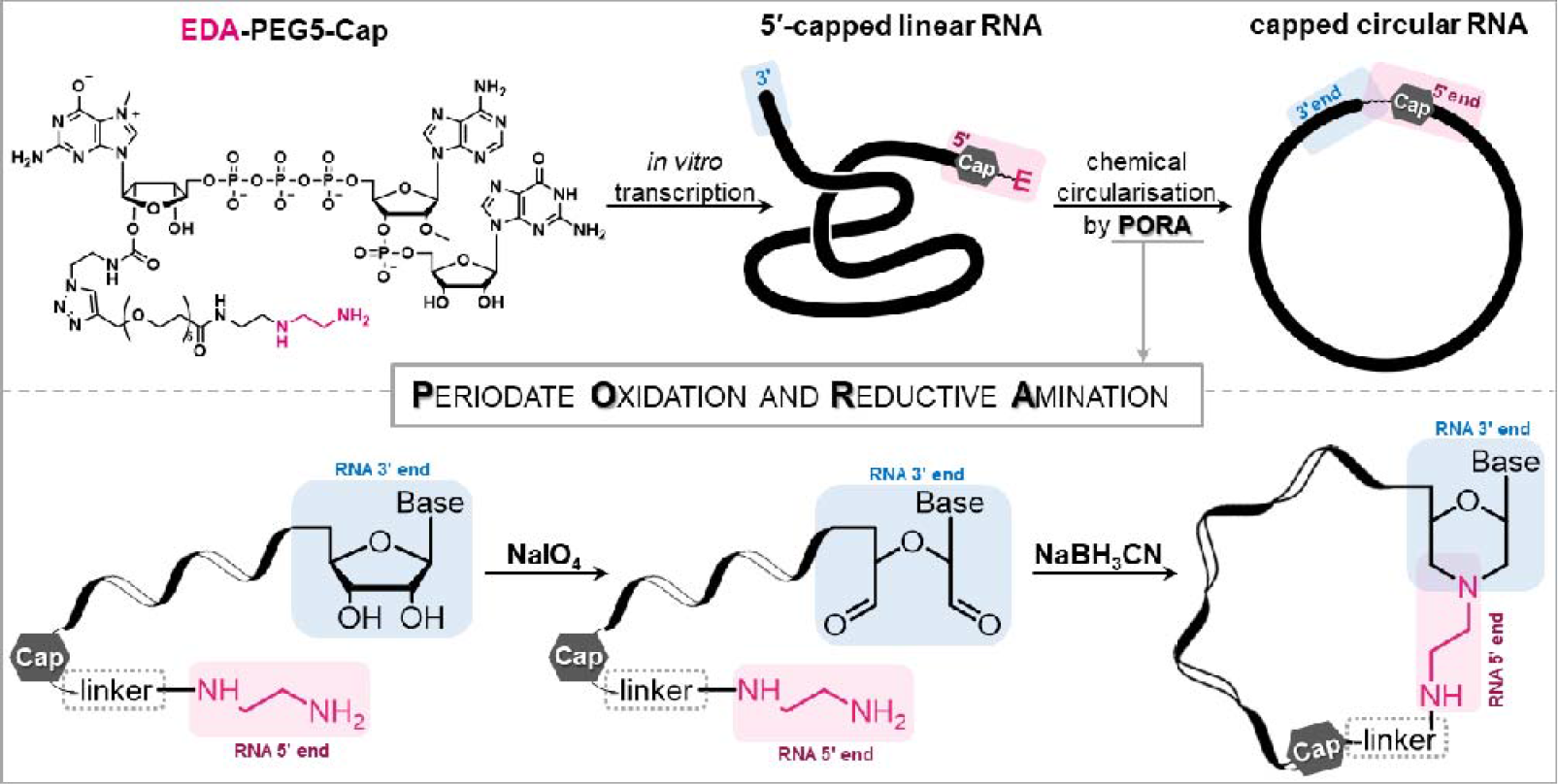

