## Supporting Information for "Chemical circularization of in vitro transcribed RNA opens new avenues for circular mRNA design"

#### Table of Contents

Figure S1 ... 2
Figure S3 ... 4
Figure S4 ... 5
Figure S5 ... 6
Figure S6 ... 7
Figure S7 ... 8
Figure S8 ... 9
Figure S9 ... 10
Figure S10 ... 11
Supplementary Table 1 ... 12
Supplementary Table 2 ... 12
Synthesis of EDA-AG ... 13
Synthesis of EDA-$PEG_5$-AG ... 13
Synthesis of $N_3$-GDP ... 14
Synthesis of $N_3$-$m^7$GDP ... 14
Synthesis of $N_3$-$m^7$Gppp$A_m$G (or $N_3$-Cap1) ... 14
Synthesis of EDA-$PEG_5$-$m^7$Gppp$A_m$G (or EDA-$PEG_5$-Cap1) ... 15
General protocol for preparation of EDA-linker-RNA ... 16
General protocol for preparation of p-RNA ... 16
Chemical circularization ... 17
Circularization with complementary oligonucleotide ... 17
Enzymatic circularization with T4 RNA Ligase I ... 18
Enzymatic circularization with T4 RNA Ligase II ... 18
RP-HPLC of circRNA ... 18
FRET probe synthesis ... 19
FRET experiments ... 19
References ... 19

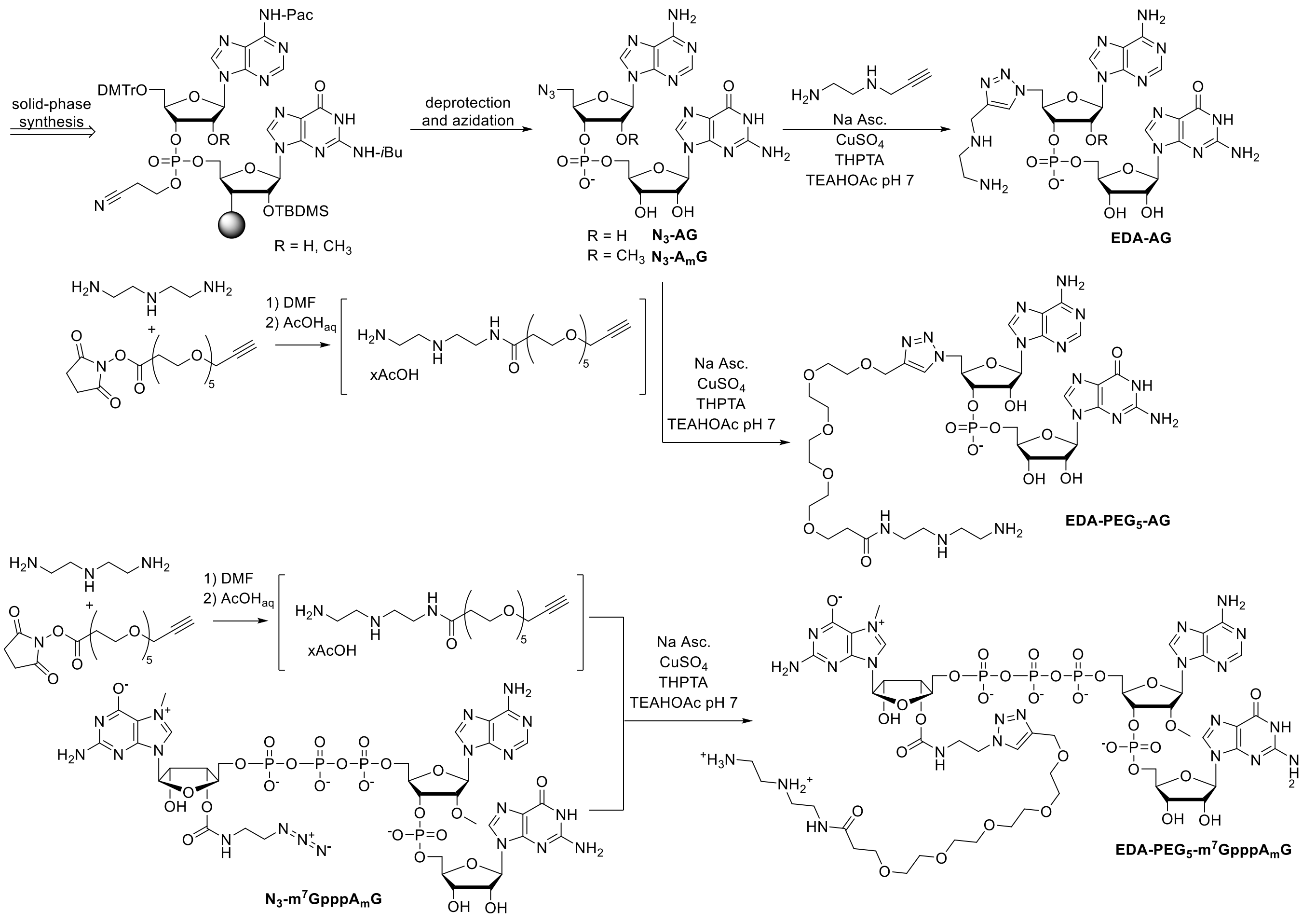


**Figure S1** Synthetic scheme for preparation of EDA-AG, EDA-PEG$_5$-AG, and EDA-PEG$_5$-m$^7$GpppA$_m$G initiators from azido-modified oligonucleotides (N$_3$-AG or N$_3$-m$^7$GpppA$_m$G).

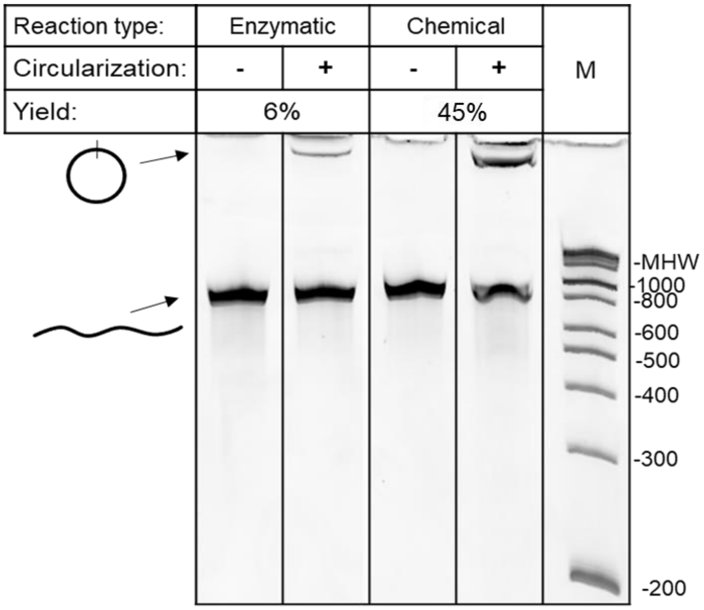


**Figure S2** Comparison of enzymatic (T4 RNA ligase I) and chemical circularization (PORA) of $RNA_{04}$.

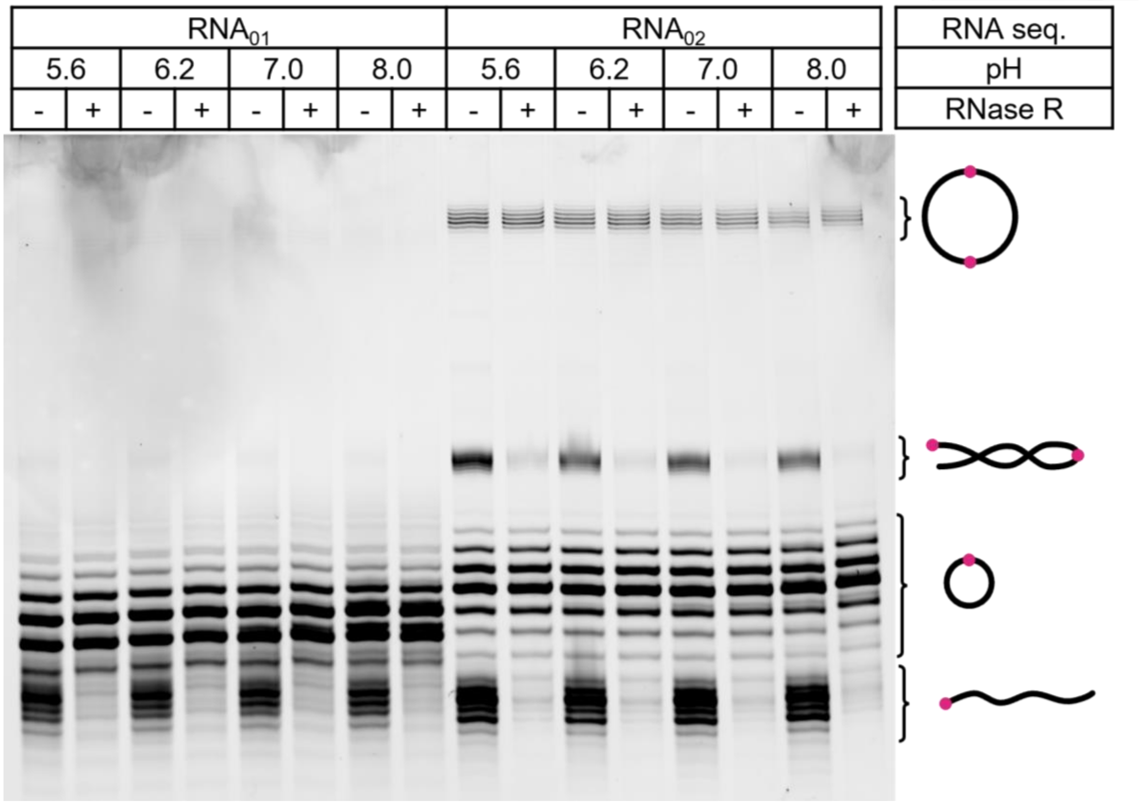


**Figure S3** Chemical circularization using two RNA oligonucleotides: E-RNA$_{01}$ (AGGGAAGCGGGCAUGCGGCCAGCC AUAGCCGAUCA) and E-RNA$_{02}$ (AGGUCAGAACGAGCGAGCGGCCAUAUGAGCAUGCA). PAGE of reaction products after performing reductive amination step at pH 5.6, 6.2, 7.0 or 8.0 (for 2 h at 20 °C) and following RNase R digestion (±RNase R). Multiple bands observed for both linear and circular RNA species stem from heterogeneity of the RNA 5' and 3' ends typical for RNAs obtained by in vitro transcription with T7RNA polymerase.

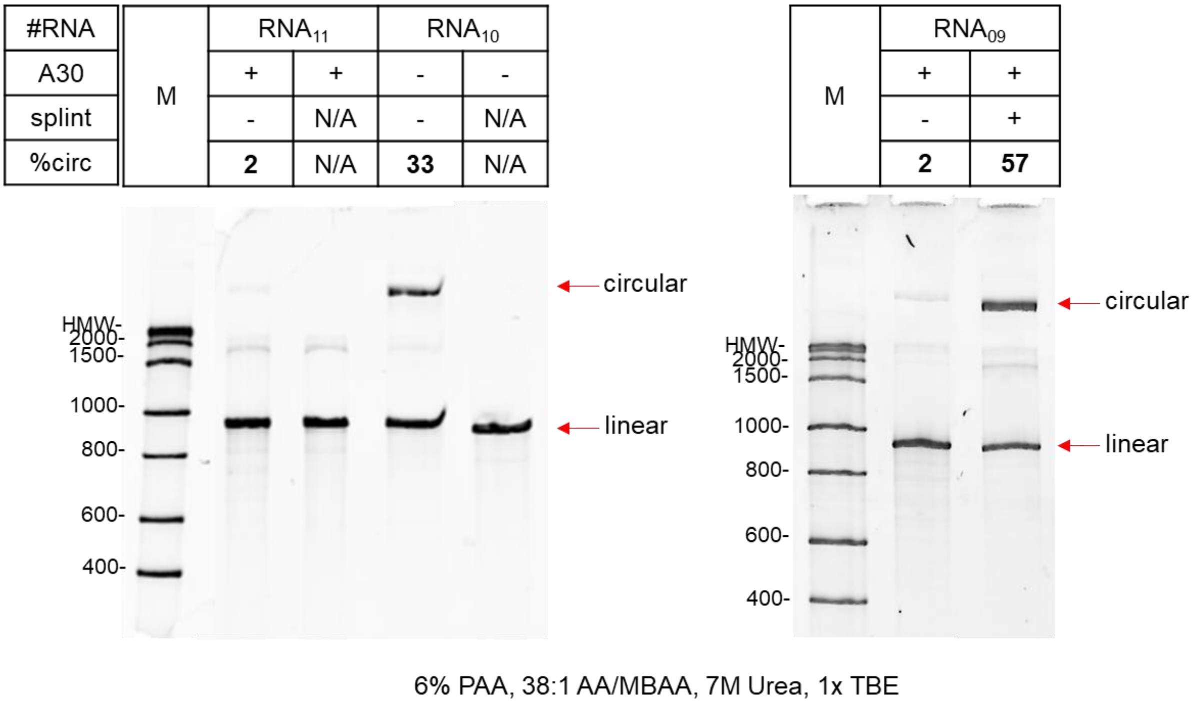


**Figure S4** Oligonucleotide splint-aided chemical circularization.

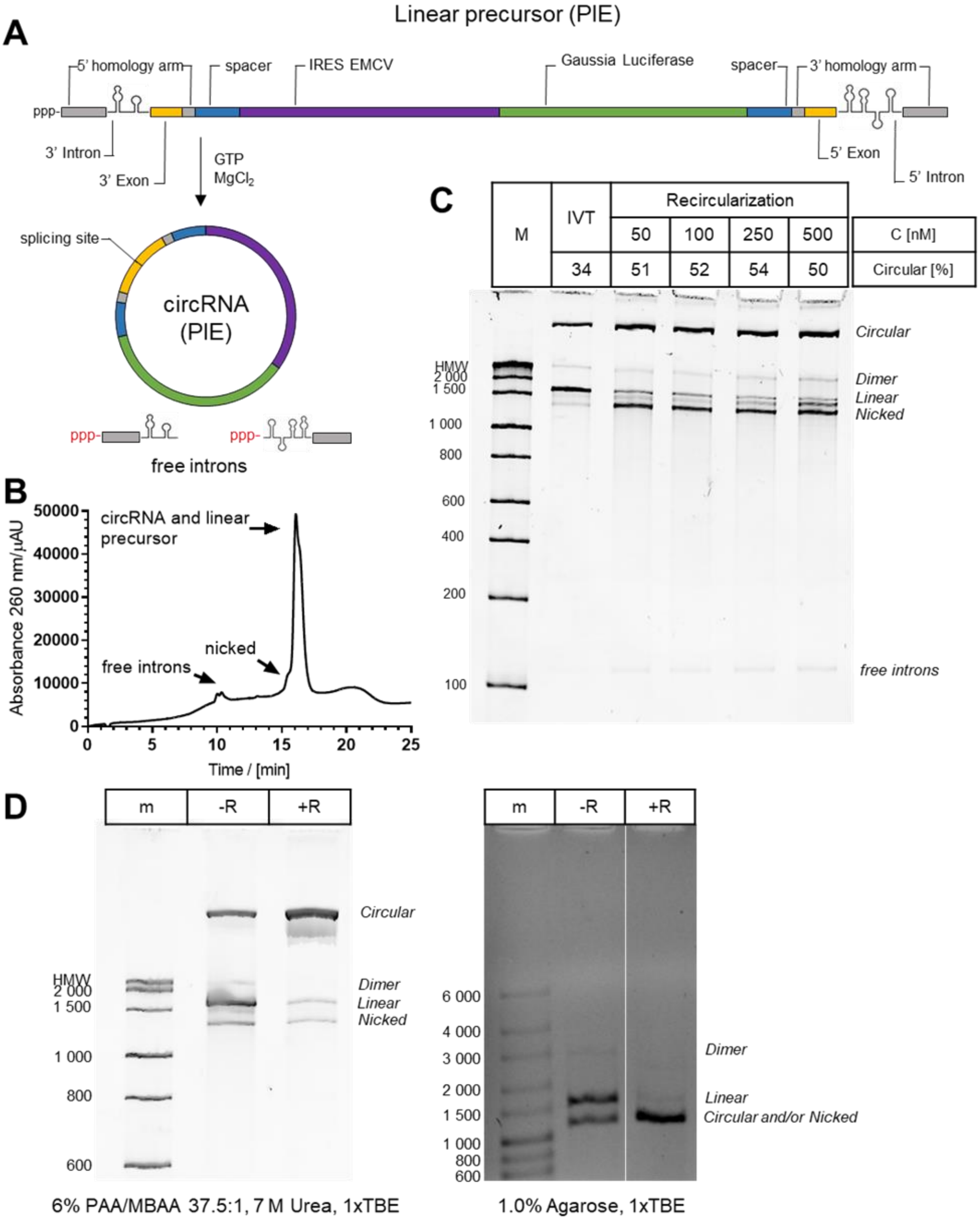


**Figure S5** Circularization of RNA using permuted intron-exon (PIE) methodology. A) General reaction scheme. B) RP-HPLC of IVT products. C) Yields of repeated self-catalytic splicing (recircularization) performed at increasing concentrations of the precursor (50–500 nM) in comparison to the IVT products. Recircularization may lead to circRNA nicking or intron elimination. D) RNase R digestion of PIE precursors monitored with polyacrylamide and agarose gel electrophoresis.

**A**

| #RNA | | $(EP_5)RNA_{13}$ | | | | | |
|---|---|---|---|---|---|---|---|
| purification | | IVT | | Cellulose | | HPLC | |
| **%circ** | M | - | **49** | - | **46** | - | **55** |
| %dimer | | - | 4 | - | 3 | - | 4 |
| %linear | | 100 | 47 | 100 | 51 | 100 | 41 |

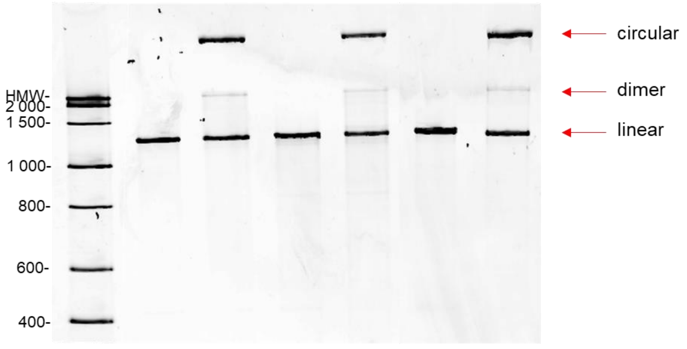


6% PAA, 38:1 AA/MBAA, 7M Urea, 1x TBE

**B**

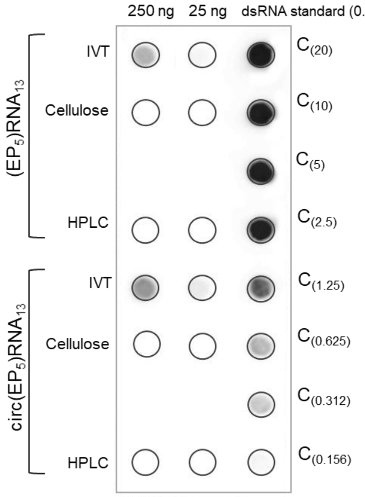


| **Sample ID** | **$(EP_5)RNA_{13}$ IVT** | **$(EP_5)RNA_{13}$ cellulose** | **$(EP_5)RNA_{13}$ HPLC** |
|---|---|---|---|
| **25 ng of RNA sample** | 6.24‰ – 12,48‰ dsRNA | n.d. <6.24‰ dsRNA | n.d. <6.24‰ dsRNA |
| **250 ng of RNA sample** | 5.00‰ – 2.50‰ dsRNA | n.d. <0.62‰ dsRNA | n.d. <0.62‰ dsRNA |

| **Sample ID** | **circ$(EP_5)RNA_{13}$ IVT** | **circ$(EP_5)RNA_{13}$ cellulose** | **circ$(EP_5)RNA_{13}$ HPLC** |
|---|---|---|---|
| **25 ng of RNA sample** | 6.24‰ – 12,48‰ dsRNA | n.d. <6.24‰ dsRNA | n.d. <6.24‰ dsRNA |
| **250 ng of RNA sample** | 2.50‰ – 5.00‰ dsRNA | n.d. <0.62‰ dsRNA | n.d. <0.62‰ dsRNA |

n.d. – dsRNA not present in the analysed sample or dsRNA content below the detection limit of method
<6.24‰ dsRNA (<0.156 ng dsRNA/25 ng of blotted mRNA)
<0.62‰ dsRNA (<0.156 ng dsRNA/250 ng of blotted mRNA)

**Figure S6** Circularization efficiency and purity of RNA [$(EP_5)RNA_{13}$] purified with different methods, followed by PAGE. A) Purification impact on circularization efficiency: Monarch® RNA Cleanup, cellulose-based chromatography and HPLC. B) Dot blot analysis of obtained RNA samples.

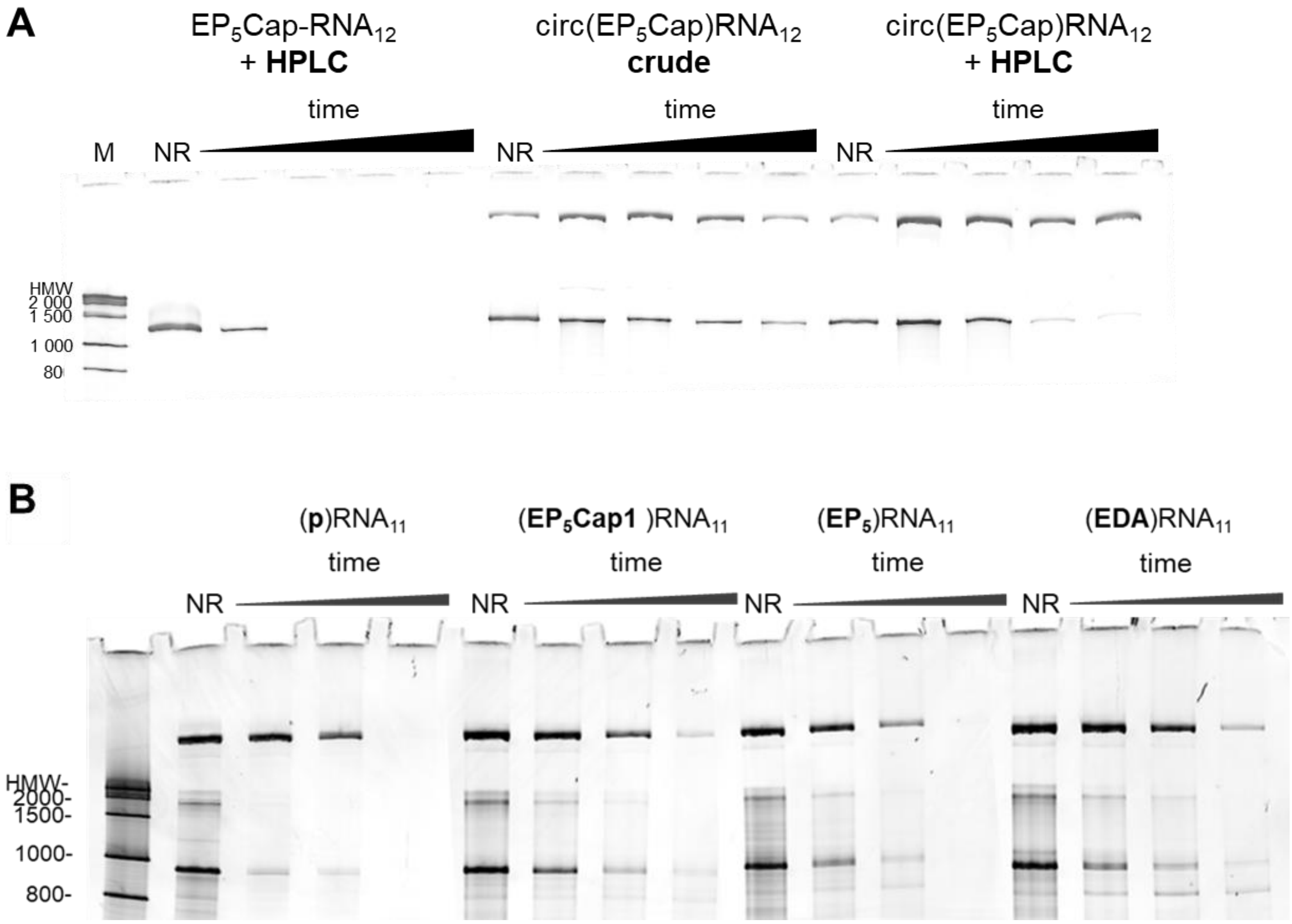


**Figure S7** Course of RNase R digestion of circRNA analogs, followed by PAGE. A) Impact of RP-HPLC purification on the course of digestion. B) Selectivity of RNase R digestion in dependence of circRNA linker length: phosphodiester bond ((p)$RNA_{11}$), EDA-$PEG_5$ motif combined with Cap1 structure (($EP_5$Cap1)$RNA_{11}$), EDA-$PEG_5$ (($EP_5$)$RNA_{11}$), and sole EDA motif ((EDA)$RNA_{11}$).

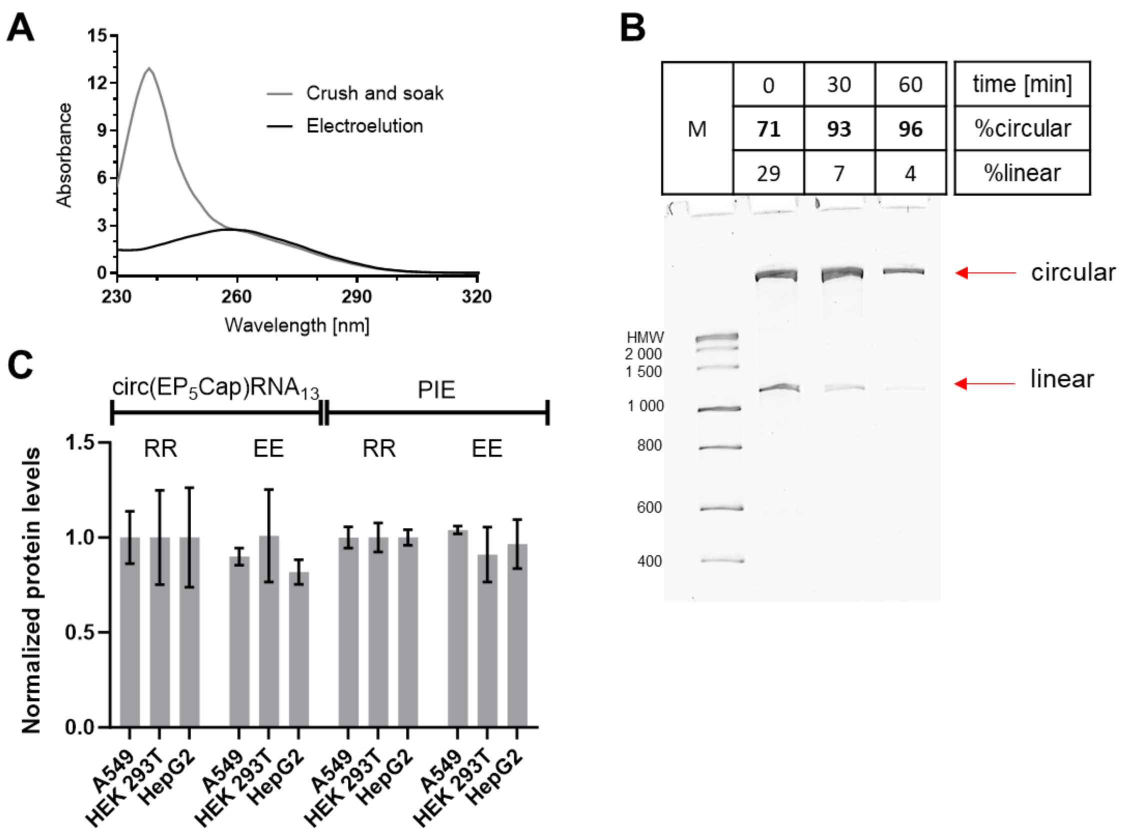


**Figure S8** A) UV spectra of the circRNA material isolated in a course of crush and soak method or electroelution. B) PAGE analysis of the circRNA material purity [% of circular in the sample] after 30 or 60 min of electroelution (0 min timepoint refer to crude material). C) Reporter protein (*Gaussia* luciferase) levels after transfection of three different cell lines (A549, HEK 293T, and HepG2) with circ($EP_5$Cap)$RNA_{13}$ or PIE $circRNA_{16}$ that were either treated with RNase R (RR) or isolated by electroelution (EE).

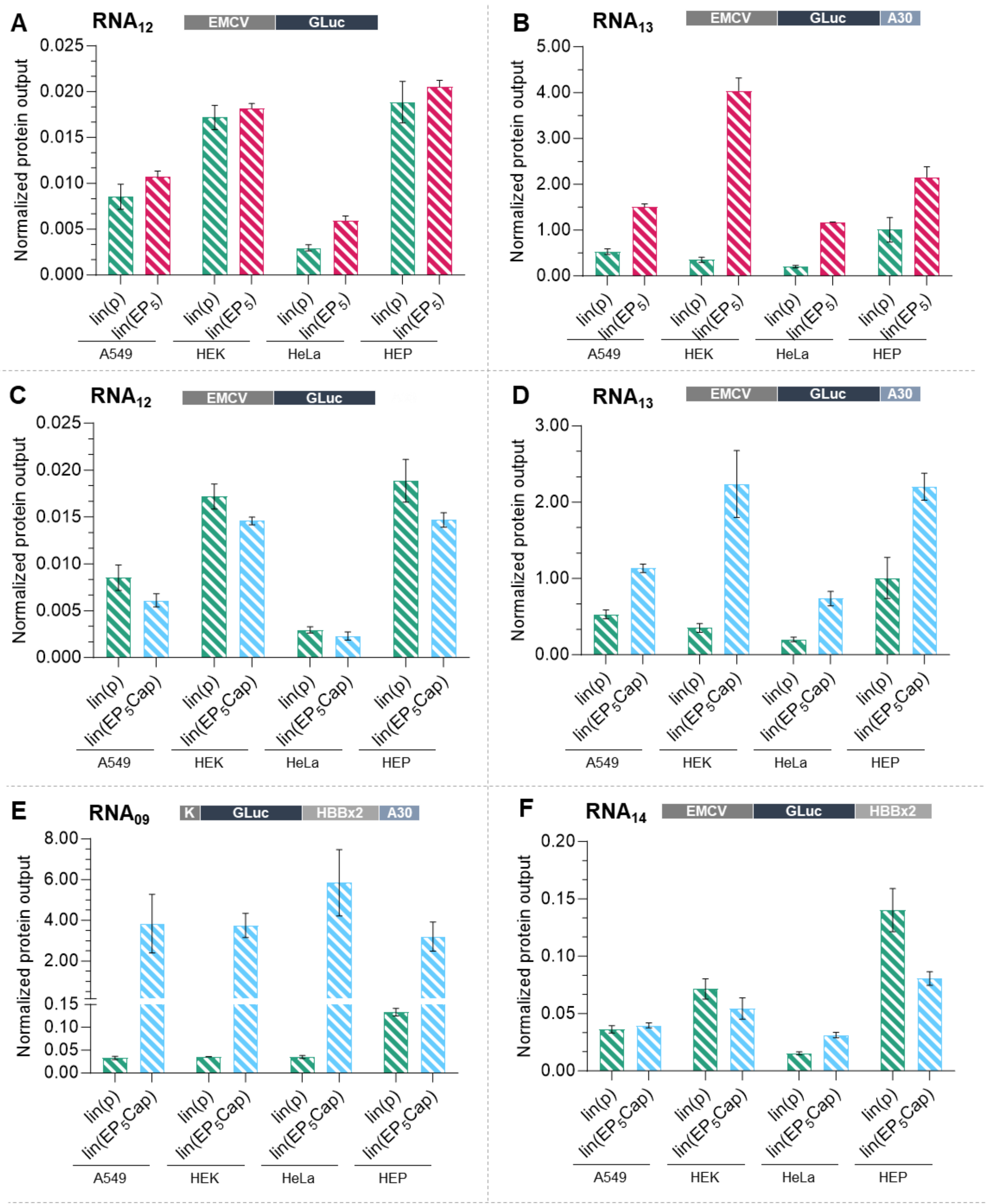


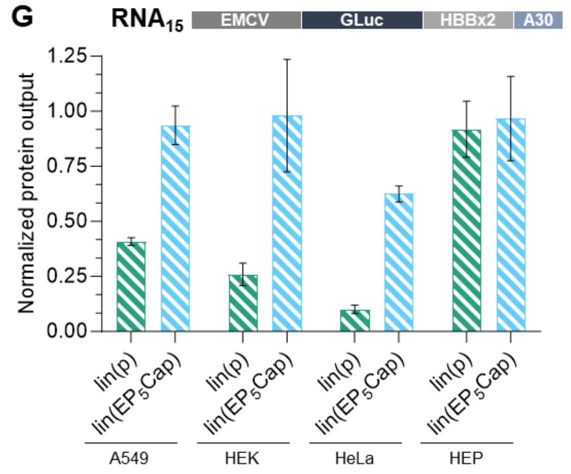


**Figure S9** Protein expression profile of linear mRNA analogues for: A) p-RNA12 and EP$_5$-RNA12; B) p-RNA13 and EP$_5$-RNA13; C) p-RNA12 and EP$_5$Cap-RNA12; D) p-RNA13 and EP$_5$Cap-RNA13; E) p-RNA09 and EP$_5$Cap-RNA09; F) p-RNA13 and EP$_5$Cap-RNA13; E) p-RNA14 and EP$_5$Cap-RNA14; G) p-RNA15 and EP$_5$Cap-RNA15. Cells of four different lines [A549, HEK293T (HEK), HepG2 (HEP), and HeLa] were transfected with linear precursors bearing 5′ monophosphate (p-), EDA-PEG$_5$ (EP$_5$-) or EDA-PEG$_5$Cap (EP$_5$Cap-) structure and coding Gaussia luciferase (RNA09, RNA12 – RNA15 sequences). The bars represent relative levels of the reporter protein harvested over 168 hours of cell culture, normalized to the level obtained for canonical mRNA reference (Cap1-RNA11). Data represents the mean and standard deviation, n=3. Comparison of the translation levels.

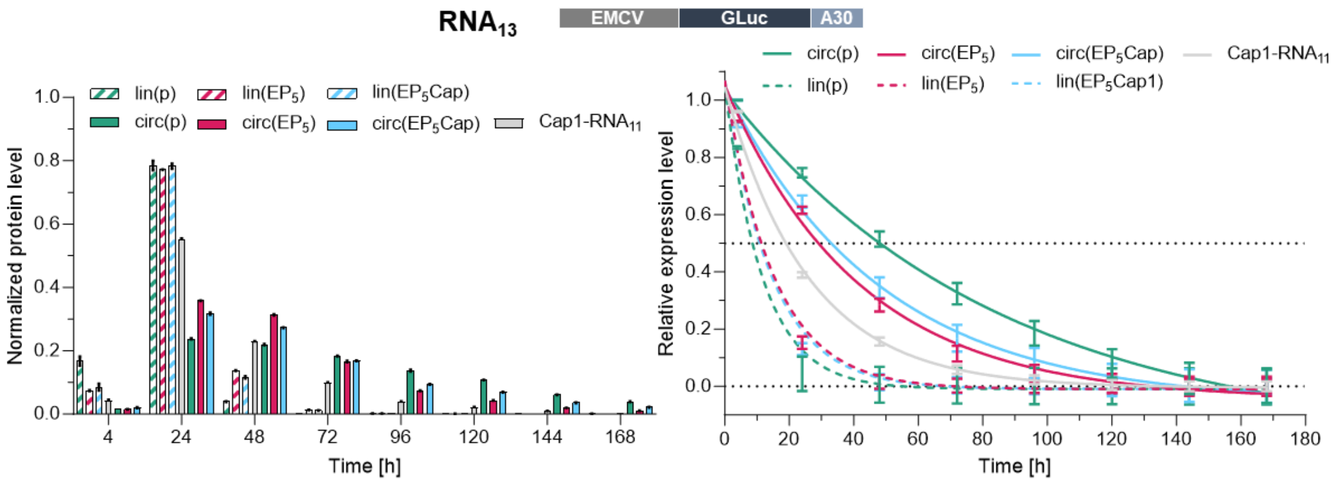


**Figure S10** Influence of the 5′-3′ linkage on the protein expression half-times in A549 cell line. The level of the reporter protein was measured for each RNA sample over 168 hours of cell culture and used to calculate the normalized protein level: a protein level at a specific timepoint divided by the total protein level collected over 168 h. The relative protein expression (one minus sum of all timepoints form 0 h up to the specific timepoint) was calculated for each timepoint and fitted to one-phase decay curve (Prism GraphPad software), that was used to determine the protein expression half-time (time after which half of the total protein level was reached).

.

**Supplementary Table 1** List of precursor RNA sequences (abbreviation, schematic of sequence elements). MCS – a multiple cloning site (84 nt); Kozak – a sequence promoting translation (8-12 nt); HBB (human beta-globin) – a sequence derived from the human beta-globin gene (50 nt); RSV – a sequence promoting $m^6A$-mediated translation (MIRES) from a respiratory syncytial virus (27 nt); EMCV – an IRES from encephalomyocarditis virus (583 nt); OligoIRES – an artificial IRES sequence (200 nt); RLuc/- – sequence of the first 62 amino acids of *Renilla* luciferase (186 nt); eGFP – sequence encoding enhanced green fluorescence protein (720 nt); GLuc – sequence coding for *Gaussia-Dura* luciferase (558 nt) HBB×2 – doubled sequence derived from human beta-globin gene (270 nt); A30 – polyadenine tail (30 nt). For full PIE construct scheme see. $E_{260}$ – molar extinction coefficient calculated for particular RNA sequence.[1]

| DNA template | Restriction enzyme | RNA sequence | RNA sequence elements | | | | RNA length | $E_{260}$ [$cm^{-1}\mu M^{-1}$] |
|---|---|---|---|---|---|---|---|---|
| | | | 5' UTR | CDS | 3' UTR | 3' tail | | |
| **DNA1** | - | $RNA_{01}$ | AGGGAAGCGGGCAUGCGGCCAGCCAUAGCCGAUCA | | | | 35 | |
| **DNA2** | - | $RNA_{02}$ | AGGUCAGAACGAGCGAGCGGCCAUAUGAGCAUGCA | | | | 35 | |
| **DNA3** | AdeI | $RNA_{03}$ | MCS | RLuc/- | - | - | 276 | 4.42 |
| **DNA4** | BamHI | $RNA_{04}$ | Kozak | eGFP | - | - | 740 | 8.93 |
| **DNA5** | BamHI | $RNA_{05}$ | Kozak | GLuc | - | - | 578 | 6.99 |
| **DNA6** | BamHI | $RNA_{06}$ | RSV | GLuc | - | - | 598 | 7.25 |
| **DNA7** | BamHI | $RNA_{07}$ | OligoIRES | GLuc | - | - | 711 | 9.35 |
| **DNA8** | BamHI | $RNA_{08}$ | HBB | GLuc | - | - | 626 | 7.58 |
| **DNA5** | AarI | $RNA_{09}$ | Kozak | GLuc | HBB ×2 | A30 | 908 | 10.99 |
| **DNA8** | PmeI | $RNA_{10}$ | HBB | GLuc | HBB ×2 | - | 918 | 10.99 |
| **DNA8** | AarI | $RNA_{11}$ | HBB | GLuc | HBB ×2 | A30 | 952 | 11.49 |
| **DNA9** | PmeI | $RNA_{12}$ | EMCV | GLuc | - | - | 1154 | 13.89 |
| **DNA9** | Esp3I | $RNA_{13}$ | EMCV | GLuc | A30 | - | 1207 | 14.39 |
| **DNA10** | PmeI | $RNA_{14}$ | EMCV | GLuc | HBB ×2 | - | 1446 | 17.24 |
| **DNA10** | Esp3I | $RNA_{15}$ | EMCV | GLuc | HBB ×2 | A30 | 1481 | 17.86 |
| **DNA11** | Eco52I | PIE | EMCV | GLuc | spacers | | 1612 | 19.61 |

**Supplementary Table 2** List of complementary DNA oligonucleotides used for RNA circularization and FRET experiments. Column descriptions: #ON – designation of a DNA oligonucleotide sequence; # RNA sequence – designation of an RNA sequence; ON sequence – sequence of the designated oligonucleotide.

| # ON | # RNA sequence | ON sequence |
|---|---|---|
| **ON1** | **$RNA_{11}$** | **TCCCTTTTTT** |
| **ON2** | | **TCTAGATTATCCCTTTTTTTTTTTTTTT** |
| **ON3** | | **GAAGCAAATGTCTAGATTATCCCTTTTTTTTTTTTTTTTTTTTTTTTT** |
| **ON4** | | **GAAGCAAATGTCTAGATTATCCCTTTTTTTTTTTTTTTTTTTTTTTTTTTTTTTTGTTTAAACATTA** |
| **ON5** | **$RNA_{09}$** | **CTTTGACTCCCATGGTGGCGTACCCCCCTTTTTTTTTTTTTTTTTTTTT** |
| **ON6** | **$RNA_{10}$** | **GAAGCAAATGTCTAGATTATCCCTTTTTTTTTTTTTTTTTTTTTTTTTTTTTTTTGTTTAAACATTA** |
| **ON7** | **$RNA_{12}$** | **AGGGGGGGAGGGAGAGGGGTACCCCCTATCCTTAGTCACCACCGGCCCCCT** |
| **ON8** | **$RNA_{13}$, $RNA_{15}$** | **GGGAGAGGGGTACCCCCTTTTTTTTTTTTTTTTTTTTT** |
| **ON9** | **$RNA_{14}$** | **GGAGGGAGAGGGGTACCCCCTAAACATTTAAATGCAATGAAAA** |

### Synthesis of EDA-AG

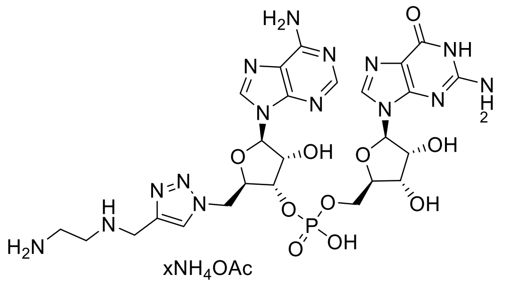


Sodium ascorbate (10.3 mg, 52 µmol) and PEDA×2HCl (3.4 mg, 20 µmol) were added to $N_3$-AG (7.4 mg, 11.5 µmol)[2] and dissolved in degassed triethylamine acetate solution (2.0 mL, 45 mM, pH 7). Then a solution of the $CuSO_4$-THPTA complex (10 µL, 100/500 mM in water) was added and incubated for 2 h at room temperature. Then an EDTA solution (20 µL, 100 mM, pH 7.0) was added and HPLC separation was performed on a C18 column flow 4.5 mL/min at 22 °C, solvent A: 100 mM $NH_4OAc$ pH 5.9; solvent B: 30% MeCN, elution: from 0 to 25% B in 40 min. After combining the fractions and freeze-drying three times, the product was obtained in the form of the ammonium acetate salt (2.8 mg, 3.50 µmol, $E_{260}$ = 24.3 $mM^{-1}cm^{-1}$) with a yield of 30%. MS ESI(-): m/z = 734.4 (Calc. $[M-H]^-$: 734.2).

### Synthesis of EDA-PEG$_5$-AG

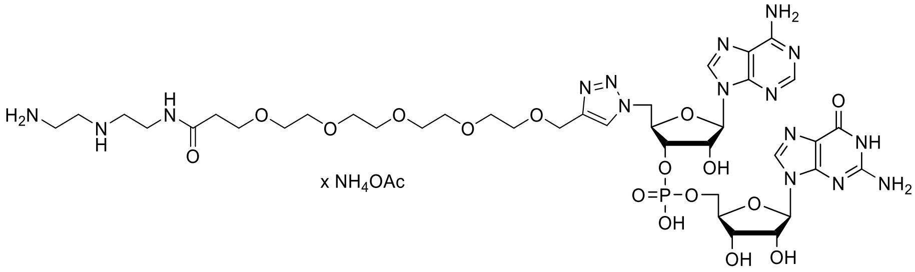


Diethylenetriamine (33.7 µl, 310 µmol, Sigma) was added to a solution of Alkyne-PEG5-NHS ester (12.5 mg, 31.2 µmol, Sigma) in DMF (200 µl). After 150 min of vortexing at 20 °C aqueous solution of ammonium acetate (50 µl, 500 mM, HPLC-grade) and acetic acid (60 µl, 50 %) were added to afford a mixture of pH ~7. Next, solution of $N_3$-AG (28 µmol, 2.00 ml)[2] and solution of Cu(II)-THPTA complex (60 µl, 50 mM) were combined with the mixture. After that, sodium ascorbate (31 mg, 157 µmol) was added, and the reaction mixture was stirred at 20 °C and monitored by HPLC. After two hours solution of triethylammonium acetate (200 µl, 1 M, HPLC-grade), solution of Cu(II)-THPTA complex (20 µl, 50 mM), and sodium ascorbate (114 mg, 586 µmol) were added, and the reaction was continued overnight. The reaction was quenched with EDTA and resolved using HPLC – column: Gemini 5 µm NX-C18 110 Å 250 × 10 mm; flow 5 ml/min at 23 °C; solvent A: 50 mM $NH_4OAc$ pH 5.9; solvent B: acetonitrile; elution: from 0% to 41% B in 25 min, then from 41% to 100% B in 10 min, then 100% B for 5 min. HPLC eluate containing the desired product was freeze-dried and resuspended in water in tree repetitions to remove the traces of solvents and ammonium acetate. The final product EDA-PEG$_5$-AG was obtained in a form of solution of an ion pair with ammonium acetate (9.2 µmol, 95% purity by HPLC) with 33% yield. MS ESI(-): m/z = 1025.8 (Calc. $[M-H]^-$: 1025.4).

### Synthesis of $N_3$-GDP

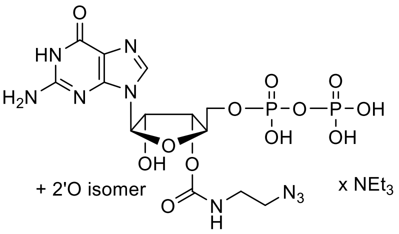


1,1-Carbonyldiimidazole (CDI) (1.21 g, 7.5 mmol, 10 eqiuv.) was added to a solution of GDP (9052.5 $mOD_{260}$, 500 mg, 750 μmol) in DMSO (15 mL). The reaction mixture was stirred at room temperature and monitored by RP-HPLC. After five hours reaction, CDI excess was hydrolyzed with water (200 μL), and the 2-azidoethylamine (7.5 mmol, 620 μL, 10 equiv.) was added, followed by DBU (3.0 mmol, 448 μL). Resulting mixture was stirred overnight at room temperature. The reaction progress was monitored by RP-HPLC. After completion of linker attachment, the solution wad was diluted with hydrochloric acid (1% aq.) until the pH 1 was reached, and the mixture was stirred overnight at room temp. The neutralized mixture was purified on DEAE-Sephadex column. The eluate was concentrated to yield a triethylammonium salt of final product as a mixture of 2′-*O*/3′*O* regioisomers (7638 $mOD_{260}$, 633 μmol, 84%).

### Synthesis of $N_3$-$m^7$GDP

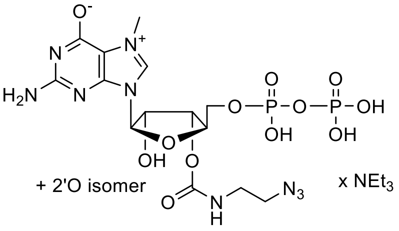


Methyl iodide (6.33 mmol, 394 μL, 10 equiv.) was added to a suspension of $N_3$-GDP (mixture of isomers) (7638 $mOD_{260}$, 350 mg, 633 μmol) in DMSO (6.33 mL), and resulting mixture was stirred at room temperature. The reaction progress was monitored by RP-HPLC. After four hours the reaction was quenched with water (60 mL) and extracted with diethyl ether (2×50 mL). The aqueous phase was neutralized with $NaHCO_3$ and purified on a DEAE-Sephadex column. The eluate was concentrated to yield a triethylammonium salt of final product as a mixture of 2′-*O*/3′*O* regioisomers (5282.6 $mOD_{260}$, 463 μmol, 82%).

### Synthesis of $N_3$-$m^7$GpppA$_m$G (or $N_3$-Cap1)

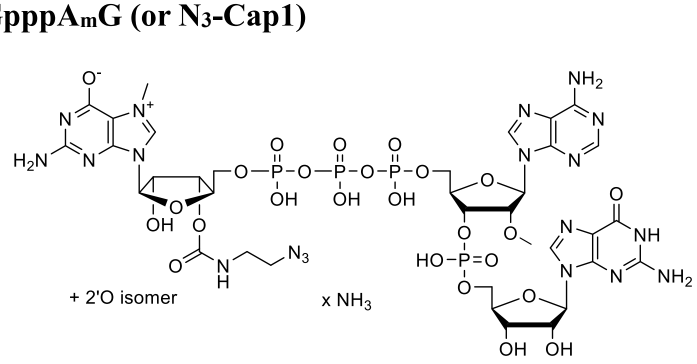


$N_3$-$m^7$GDP (mix of regioisomers) (1778 $mOD_{260}$, 156 μmol, 88 mg, 1.5 equiv) and Im-pA$_m$pG (2833.6 $mOD_{260}$, 78.5 mg, 104 μmol)[3] were suspended in DMF (2.0 mL) and $ZnCl_2$ (228 mg, 1.67 mmol, 16 equiv.)

was added. Resulting mixture was stirred, and the reaction progress was monitored by RP-HPLC. After completion, reaction was terminated by addition of aqueous EDTA (20 mL), and the pH was adjusted to 6 with solid $NaHCO_3$. The product was purified by semipreparative RP-HPLC with use of linear gradient (0-40%) of acetonitrile in ammonium acetate (pH 5.9, 0.05 M) over 60 min. Collected fractions were combined and concentrated to remove the excess of ACN. The product (mixture of 2′-*O*/3′*O* regioisomers) (3232 $mOD_{260}$, 55 µmol, 53%) was obtained as a white powder after lyophilization. $^1$H NMR (500 MHz, $D_2O$, 25 °C): δ = 9.21 (s, 1H, $H8_{m7G}$, isomer 3′), 9.17 (s, 1H, $H8_{m7G}$, isomer 2′) 8.57 (s, 2H, $H8_A$, isomer 2′ and 3′), 8.31 (s, 2H, $H2_A$, isomer 2′ and 3′), 8.04 (s, 2H, $H8_G$, isomer 2′ and 3′), 6.10 (m, 2H, $H1'_A$, isomer 2′ and 3′), 6.02 (d, $^3J_{H\text{-}H}$ = 3.3 Hz, 1H, $H1'_{m7G}$ isomer 2′), 5.89 (d, $^3J_{H\text{-}H}$ = 5.6 Hz, 1H, $H1'_{m7G}$ isomer 3′), 5.80 (m, 2H, $H1'_G$, isomer 2′ and 3′), 5.47 (m, 1H, $H2'_{m7G}$ isomer 2′), 5.23 (m, 1H, $H3'_{m7G}$ isomer 3′), 4.93 (m, 2H, $H3'_A$, isomer 2′ and 3′), 4.78 (m, overlapped with HDO, 2H, $H2'_G$, $H2'_{m7G}$ isomer 3′), 4.67 (m, 1H, $H3'_{m7G}$ isomer 2′), 4.54–4.46 (m, 2H, $H3'_G$, $H4'_{m7G}$ isomer 3′, 2H, $H4'_A$, isomer 2′ and 3′), 4.45–4.41 (m, 2H, $H2'_A$, isomer 2′ and 3′), 4.35 (m, 2H, $H4'_G$, $H4'_{m7G}$ isomer 2′, 4H, $H5'_{m7G}$, $H5''_{m7G}$, isomer 2′ and 3′), 4.30–4.14 (m, 8H, $H5'_G$, $H5''_G$, $H5'_A$, $H5''_A$, isomer 2′ and 3′), 4.04 (s, 6H *N*7-$CH_3$ $_{m7G}$, isomer 2′ and 3′), 3.49 (m, 6H, $CH_3$, 2′-O-$CH_3$, isomer 2′ and 3′), 3.45–3.30 (m, 8H, $CH_2$-$CH_2$, isomer 2′ and 3′).

#### Synthesis of EDA-PEG$_5$-m$^7$GpppA$_m$G (or EDA-PEG$_5$-Cap1)

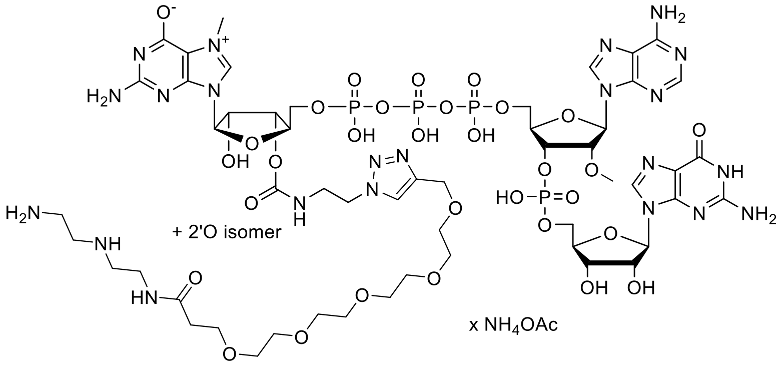


Diethylenetriamine (9.6 µl, 90 µmol, Sigma) was added to a solution of Alkyne-PEG5-NHS ester (12.0 mg, 29.9 µmol, Sigma) in DMF (180 µl). After 60 min of vortexing at 25 °C aqueous solution of triethylammonium acetate (520 µl, 50 mM, HPLC-grade) and acetic acid (62 µl, 50 %) were added to afford a mixture of pH 6-7. Next, solution of $N_3$-m$^7$GpppA$_m$G (400 µl, 26 µmol) and solution of Cu(II)-THPTA complex (96 µl, 50 mM) were combined with the mixture. After that, sodium ascorbate (49 mg, 248 µmol) was added, and the reaction mixture was stirred at 25 °C and monitored by HPLC. After two hours solution of Cu(II)-THPTA complex (48 µl, 50 mM), and sodium ascorbate (25 mg, 126 µmol) were added, and the reaction was continued overnight. The reaction was quenched with EDTA and resolved using HPLC – column: Gemini 5 µm NX-C18 110 Å 250 × 10 mm; flow 5 ml/min at 23 °C; solvent A: 50 mM $NH_4OAc$ pH 5.9; solvent B: water/acetonitrile mixture 1:2 (v/v); elution: from 0% to 25% B in 45 min, then from 25% to 100% B in 5 min, then 100% B for 5 min. HPLC eluate containing the desired product was freeze-dried and resuspended in water in tree repetitions to remove the traces of solvents and ammonium acetate. The final product EDA-PEG$_5$-AG was obtained in a form of solution of an ion pair with ammonium acetate (10.5 µmol, 99% purity by HPLC) with 40% yield. HRMS: m/z = 1631.42458, 815.20983 (calc. $[M\text{-}H]^-$: 1631.42375, $[M\text{-}2H]^{2-}$: 815.20824).

**DNA template preparation**

To afford the desired DNA template for IVT plasmids presented in supplementary table were linearized with a restriction enzyme according to manufacturer’s protocol (Thermo). For details see Supplementary Table 1.

### General protocol for preparation of EDA-linker-RNA

| | $C_{stock}$ | $V_{stock}$ [µl] | $C_{fin}$ |
|---|---|---|---|
| $H_2O$ | | up to 50 µl | |
| Transcription Buffer ×5 | ×5 | 10 | ×1 |
| U/A/CTP [mM] | 33.3 | 9 | 6 |
| GTP [mM] | 100 | 2 | 4 |
| EDA-linker-AG (or EDA-linker-Cap) [mM] | 40 | 15 | 12 |
| DNA template [ng/µl] | 400 | 5 | 40 |
| $MgCl_2$ [mM] | 1 000 | 1.25 | 25 |
| Ppase [U/µl] | 0.1 | 1 | 0.002 |
| Ribolock [U/µl] | 40 | 1.25 | 1 |
| T7 RNAP HC [U/µl] | 200 | 5 | 20 |

All ingredients of the reaction mixture listed in the table above were mixed in room temperature in the following order: water, Transcription Buffer ×5 (200 mM Tris-HCl pH 7.9, 30 mM $MgCl_2$, 50 mM DTT, 50 mM NaCl and 10 mM spermidine, Thermo), NTPs (Thermo), EDA-linker-AG (or EDA-linker-Cap), DNA template, $MgCl_2$, Ribolock (Thermo), and T7 RNA polymerase HC (Thermo). The reaction mixture was vortexed, gently centrifuged, and incubated for 2 h at 37 °C. Next, DNase I (2.0 µl, 1 U/µl, Thermo) was added, and the mixture was incubated for 30 min at 37 °C. The RNA product was isolated form reaction mixture using Monarch RNA cleanup kit (500 µg, NEB). The isolated RNA (~200 µg) was resolved using RP-HPLC – column: PolymerX RP-1 3 µm 100 Å, 4.1 × 150 mm; flow 1.0 ml/min at 60 °C; solvent A: 100 mM TEAA pH 7.0; solvent B: 200 mM TEAA pH 7.0 / acetonitrile 1:1 (vol/vol); elution: from 20% to 32.5% B in 25 min, then from 32.5% to 100% B in 2 min, then 100% B for 2 min. HPLC eluate containing the desired RNA product was concentrated by precipitation: sodium acetate (0.1 V, 3.0 M, pH 5.2), and isopropanol (1.0-1.5 V) were added to the eluate (initial volume = 1 V), and the solution was cooled for 30 min at -80 °C or overnight at -20 °C. The pellet was centrifuged (30 min, 14 000 g, 4 °C), washed with 80% ethanol (1.0 ml), centrifuged (10 min, 14 000 g, 4 °C), dried *in vacuo*, and suspended in water to afford the RNA product (100-180 µg, 50-90% of the crude product).

### General protocol for preparation of p-RNA

| | $C_{stock}$ | $V_{stock}$ [µl] | $C_{fin}$ |
|---|---|---|---|
| $H_2O$ | | up to 50 µl | |
| Transcription Buffer ×5 | ×5 | 10 | ×1 |
| U/A/CTP [mM] | 33.3 | 9 | 6 |
| GTP [mM] | 100 | 1.5 | 3 |
| GMP [mM] | 100 | 10 | 20 |
| DNA template [ng/µl] | 400 | 5 | 40 |
| $MgCl_2$ [mM] | 1 000 | 1.25 | 25 |
| Ppase [U/µl] | 0.1 | 1 | 0.002 |
| Ribolock [U/µl] | 40 | 1.25 | 1 |
| T7 RNAP HC [U/µl] | 200 | 5 | 20 |

All ingredients of the reaction mixture listed in the table above were mixed at 37 °C in the following order: water, Transcription Buffer ×5 (200 mM Tris-HCl pH 7.9, 30 mM $MgCl_2$, 50 mM DTT, 50 mM NaCl and 10 mM spermidine, Thermo), NTPs (Thermo), GMP (preincubated at 37 °C), DNA template, $MgCl_2$,

Ribolock (Thermo), and T7 RNA polymerase HC (Thermo). The reaction mixture was vortexed, gently centrifuged, and incubated for 2 h at 37 °C. Next, DNase I (2.0 µl, 1 U/µl, Thermo) was added, and the mixture was incubated for 30 min at 37 °C. The crude RNA product was isolated form reaction mixture using Monarch RNA cleanup kit (500 µg, NEB). The isolated crude RNA (~150 µg, 2.5-3.5 g of RNA per one litter of IVT reaction mixture) was resolved using RP-HPLC-column: PolymerX RP-1 3 µm 100 Å, 4.1 × 150 mm; flow 1.0 ml/min at 60 °C; solvent A: 100 mM TEAA pH 7.0; solvent B: 200 mM TEAA pH 7.0 / acetonitrile 1:1 (vol/vol); elution: from 20% to 32.5% B in 25 min, then from 32.5% to 100% B in 2 min, then 100% B for 2 min. HPLC eluate containing the desired RNA product was concentrated by precipitation: sodium acetate (0.1 V, 3.0 M, pH 5.2), and isopropanol (1.0-1.5 V) were added to the eluate (initial volume = 1 V), and the solution was cooled for 30 min at -80 °C or overnight at -20 °C. The pellet was centrifuged (30 min, 14 000 g, 4 °C), washed with 80 % ethanol (1.0 ml), centrifuged (10 min, 14 000 g, 4 °C), dried *in vacuo*, and suspended in water to afford the RNA product (70-130 µg, 50-90% of the crude product).

### Chemical circularization

A diluted solution of the precursor RNA (30-40 µg in 490 µl, ~140 µM, pre-circRNA 2, 4-6, 8, 10, 12, 13, 15, 17) was mixed with buffer (70 µl, 800 mM $KH_2PO_4$, 200 mM NaCl, pH 7.0) and incubated at 65 °C for 5 min. After that, hot solution was transferred on bench, where it cooled to room temperature. After 10-15 min at room temperature, fresh solution of sodium periodate (70 µl, 10 mM) was added, and the reaction mixture was immediately placed in dark and incubated at 25 °C. After 30 min, fresh solution of sodium cyanoborohydride (70 µl, 200 mM) was added, and the incubation was continued for next 90 min. The crude circRNA product was isolated form reaction mixture using Monarch RNA cleanup kit (50 µg, NEB). The isolated RNA (80-99% of the initial amount) was resolved using RP-HPLC-column: PolymerX RP-1 3 µm 100 Å, 4.1 × 150 mm; flow 1.0 ml/min at 60 °C; solvent A: 100 mM HAA pH 7.0; solvent B: 100 mM HAA pH 7 in 75 % acetonitrile; elution: from 55% to 60% B in 25 min, then from 60% to 100% B in 2 min, then 100% B for 2 min. HPLC eluate containing the desired RNA product was concentrated by precipitation: sodium acetate (0.1 V, 3.0 M, pH 5.2), and isopropanol (1.0-1.5 V) were added to the eluate (initial volume = 1 V), and the solution was cooled for 30 min at -80 °C or overnight at -20 °C. The pellet was centrifuged (30 min, 14 000 g, 4 °C), washed with 80% ethanol (1.0 ml), centrifuged (10 min, 14 000 g, 4 °C), dried *in vacuo*, and suspended in water to afford the circRNA product (5-10 µg, 15-25% of the initial amount). For additional purification and removal of linear RNA, the product was treated with RNase R as described below (see section RNase R digestion).

### Circularization with complementary oligonucleotide

A diluted solution of the precursor RNA (30-40 µg in 480 µl, ~145 µM, pre-circRNA 2, 4-6, 8, 10, 12, 13, 15, 17) was mixed with buffer (70 µl, 800 mM $KH_2PO_4$, 200 mM NaCl, pH 7.0), and complementary oligonucleotide (~11 µl, 10 µM, 1.5 eq, see Supplementary Table 2 for sequence) and incubated at 65 °C for 5 min. After that, hot solution was transferred on bench, where it cooled to room temperature. After 10-15 min at room temperature, fresh solution of sodium periodate (70 µl, 10 mM) was added, and the reaction mixture was immediately placed in dark and incubated at 25 °C. After 30 min, fresh solution of sodium cyanoborohydride (70 µl, 200 mM) was added, and the incubation was continued for next 90 min. The crude circRNA product was isolated form reaction mixture using Monarch RNA cleanup kit (50 µg, NEB). The isolated RNA was resolved using RP-HPLC-column: PolymerX RP-1 3 µm 100 Å, 4.1 × 150 mm; flow 1.0 ml/min at 60 °C; solvent A: 100 mM HAA pH 7.0; solvent B: 100 mM HAA pH 7 in 75% acetonitrile; elution: from 55% to 60% B in 25 min, then from 60% to 100% B in 2 min, then 100% B for 2 min. HPLC eluate containing the desired RNA product was concentrated by precipitation: sodium acetate (0.1 V, 3.0 M, pH 5.2), and isopropanol (1.0-1.5 V) were added to the eluate (initial volume = 1 V), and the solution was cooled for 30 min at -80 °C or overnight at -20 °C. The pellet was centrifuged (30 min, 14 000 g, 4 °C), washed with 80% ethanol (1.0 ml), centrifuged (10 min, 14 000 g, 4 °C), dried *in vacuo*, and suspended in water to afford the circRNA product (5-10 µg, 15-25% of the initial amount). For additional purification and removal of linear RNA, the product was treated with RNase R as described below (see section RNase R digestion).

### Enzymatic circularization with T4 RNA Ligase I

A diluted solution of the precursor RNA (10-15 µg in 95 µl, ~150 µM, pre-circRNA 1, 3, 7, 9, 11, 14, 16) was mixed with buffer (5 µl, 100 mM Tris-HCl, 500 mM NaCl, pH 7.0) and incubated at 65 °C for 5 min. After that, hot solution was transferred on bench, where it cooled to room temperature. After 10-15 min at room temperature, T4 RNA ligase 1 buffer ×10 (15 µl, 500 mM Tris-HCl, 100 mM $MgCl_2$, 10 mM DTT, pH 7.5, NEB), PEG 8000 (15 µl, 50%, NEB), ATP (0.75 µl, 10 mM, NEB), RiboLock RNase inhibitor (3.75 µl, 40 U/µl, Thermo), and T4 RNA ligase 1 (15 µl, 10 U/µl, NEB) were added, and the reaction mixture was incubated at 25 °C for 150 min. The crude circRNA product was isolated form reaction mixture using Monarch RNA cleanup kit (10 µg, NEB). The isolated RNA (80-99% of the initial amount) was resolved using RP-HPLC-column: RNASept Prep 2 µm 100 Å, 7.8 × 50 mm; flow 1.0 ml/min at 60 °C; solvent A: 100 mM HAA pH 7.0 in 10% acetonitrile; solvent B: 100 mM HAA pH 7 in 75% acetonitrile; elution: from 20% to 70% B in 25 min, then from 70% to 100% B in 2 min, then 100% B for 2 min. HPLC eluate containing the desired RNA product was concentrated by precipitation: sodium acetate (0.1 V, 3.0 M, pH 5.2), and isopropanol (1.0-1.5 V) were added to the eluate (initial volume = 1 V), and the solution was cooled for 30 min at -80 °C or overnight at -20 °C. The pellet was centrifuged (30 min, 14 000 g, 4 °C), washed with 80% ethanol (1.0 ml), centrifuged (10 min, 14 000 g, 4 °C), dried *in vacuo*, and suspended in water to afford the circRNA product (1-4 µg, 10-30% of the crude product). For additional purification and removal of linear RNA, the product was treated with RNase R as described below (see section RNase R digestion).

### Enzymatic circularization with T4 RNA Ligase II

A diluted solution of the precursor RNA (30-40 µg in 220 µl, ~320 µM, pre-circRNA 1, 3, 7, 9, 11, 14, 16) was mixed with complementary oligonucleotide (~11 µl, 10 µM, 1.5 eq, see Supplementary Table 2 for sequence) and incubated at 65 °C for 5 min. The hot solution was then transferred on bench, where it cooled to room temperature. After 10-15 min at room temperature, T4 RNA ligase 2 buffer ×10 (35 µl, 500 mM Tris-HCl, 20 mM $MgCl_2$, 10 mM DTT, 4 mM ATP, pH 7.5, NEB), PEG 8000 (70 µl, 50%, NEB), RiboLock RNase inhibitor (8.75 µl, 40 U/µl, Thermo), and T4 RNA ligase 2 (3.5 µl, 10 U/µl, NEB) were added, and the reaction mixture was incubated at 25 °C for 150 min. The crude circRNA product was isolated form reaction mixture using Monarch RNA cleanup kit (500 µg, NEB). The isolated RNA (80-99% of the initial amount) was resolved using RP-HPLC-column PolymerX RP-1 3 µm 100 Å, 4.1 × 150 mm; flow 1.0 ml/min at 60 °C; solvent A: 100 mM HAA pH 7.0; solvent B: 100 mM HAA pH 7 in 75% acetonitrile; elution: from 55% to 60% B in 25 min, then from 60% to 100% B in 2 min, then 100% B for 2 min. HPLC eluate containing the desired RNA product was concentrated by precipitation: sodium acetate (0.1 V, 3.0 M, pH 5.2), and isopropanol (1.0-1.5 V) were added to the eluate (initial volume = 1 V), and the solution was cooled for 30 min at -80 °C or overnight at -20 °C. The pellet was centrifuged (30 min, 14 000 g, 4 °C), washed with 80% ethanol (1.0 ml), centrifuged (10 min, 14 000 g, 4 °C), dried *in vacuo*, and suspended in water to afford the circRNA product (5-10 µg, 15-25% of the initial amount). For additional purification and removal of linear RNA, the product was treated with RNase R as described below (see section RNase R digestion).

### RP-HPLC of circRNA

All HPLC separations were performed on modular, low-pressure gradient HPLC apparatus manufactured by Shimadzu (LC-40/LC-20 series) with thermostated column holder or column oven, DAD detector and fluorescence detector. Mobile phase was composed of non-autoclaved MQ water, HPLC-grade acetonitrile (J.T.Baker), high quality triethylamine (Bioultra, Sigma), HPLC-grade n-hexylamine (Ottokemi), and

HPLC-grade glacial acetic acid (J.T.Baker). Solvents of desired composition, concentration and pH were filtered with 0.45 μm filter and stored in dark at 4 °C for longer periods of time. Chromatograms were recorded with detection of absorbance at 260 nm (ref 360 nm, 10 nm bandwidth), absorbance spectrum (220-700 nm), and fluorescence of interest (Cy3 – 550/565, Cy5- 650/665).

### FRET probe synthesis

The Cy5-RNA1-Cy3 FRET probe was synthesized according to previously reported protocol.[2] First in vitro transcription of RNA1 sequence was performed in presence of $N_3$-AG transcription initiator. The crude product of IVT was isolated by Monarch RNA cleanup kit (NEB) and subjected to 5′-Cy5 and 3′-Cy3 dual labeling. The crude labeling product was isolated from labeling reaction mixture by Monarch RNA cleanup kit (NEB) and subsequently resolved with HPLC. The HPLC eluate containing the desired product was concentrated by precipitation dried *in vacuo* and suspended in water to afford the Cy5-RNA1-Cy3 FRET probe solution.

### FRET experiments

The Cy5-RNA1-Cy3 FRET probe solution was diluted with buffer (80 mM $KH_2PO_4$, 20 mM NaCl, pH 7.0, final volume of 100 μl) to afford final probe concentration of 100 nM and transferred to a quartz fluorescence cuvette (1 × 1 × 350 mm). Optionally, the solution of a complementary DNA oligonucleotide (2 μl, 10 μM, ON2-ON5) was added. The mixture was heated to from 20 to 90 °C in 8 min and gradually cooled to 20 °C or 4 °C in 20-30 min. Emission spectra were recorded on a Cary Eclipse spectrofluorometer (Agilent), equipped with a xenon lamp, single cell peltier accessory (spvf 1×0, Agilent), excitation at 500 nm, emission 510-800 nm, 10 mm slit.
